# Sphingomyelin suppresses Hedgehog signaling by restricting cholesterol accessibility at the ciliary membrane

**DOI:** 10.1101/699819

**Authors:** Maia Kinnebrew, Ellen J. Iverson, Bhaven B. Patel, Ganesh V. Pusapati, Jennifer H. Kong, Kristen A. Johnson, Giovanni Luchetti, Douglas F. Covey, Christian Siebold, Arun Radhakrishnan, Rajat Rohatgi

## Abstract

Transmission of the Hedgehog signal across the plasma membrane by Smoothened is proposed to be triggered by its direct interaction with cholesterol. But how is cholesterol, an abundant lipid, regulated tightly enough to control a signaling system that can cause birth defects and cancer? Using toxin-based sensors that distinguish between distinct pools of cholesterol, we find here that Smoothened activation and Hedgehog signaling are driven by a biochemically defined fraction of membrane cholesterol, termed accessible cholesterol. Increasing accessible cholesterol levels by depletion of sphingomyelin, which sequesters cholesterol in complexes, potentiates Hedgehog signaling. By inactivating the transporter-like protein Patched 1, Hedgehog ligands trigger an increase in cholesterol accessibility in the ciliary membrane, the subcellular location for Smoothened signaling. Thus, compartmentalization of Hedgehog signaling in the primary cilium may allow cholesterol accessibility to be used as a second messenger to mediate the communication between Patched 1 and Smoothened, without causing collateral effects on other cellular processes.

## Introduction

A long-standing mystery in Hedgehog (HH) signaling is how Patched 1 (PTCH1), the receptor for HH ligands, inhibits Smoothened (SMO), a G-protein-coupled receptor (GPCR) family protein that transduces the HH signal across the membrane (Kong, Siebold, and Rohatgi 2019). The observation that cholesterol can directly bind and activate SMO has led to the proposal that PTCH1 regulates SMO by restricting its access to cholesterol (E. F. Byrne et al. 2016; Luchetti et al. 2016; P. Huang et al. 2016). Structural and biochemical studies have confirmed that PTCH1 could be a cholesterol transporter; however, transport activity has not yet been demonstrated either in a purified system or at endogenous expression levels in cells (Zhang et al. 2018; Gong et al. 2018; X. Qi et al. 2018; Bidet et al. 2011; C. Qi et al. 2018; Qian et al. 2019). Furthermore, the resolution of the PTCH1 cryo-EM structures is not high enough to distinguish cholesterol from other sterol lipids as PTCH1 substrates.

A challenge to this model is presented by the fact that cholesterol constitutes up to 50% of the lipid molecules in the plasma membrane (Y. Lange et al. 1989; A. Das et al. 2013; Touster et al. 1970; Colbeau, Nachbaur, and Vignais 1971): how can such an abundant lipid be kept away from SMO to prevent inappropriate pathway activation? Indeed, other less abundant lipids can bind and regulate SMO activity, including oxysterols, phosphoinositides, endocannabinoids and arachidonic acid derivatives (Nachtergaele et al. 2012; Khaliullina et al. 2015; Arensdorf et al. 2017; Jiang et al. 2016). Side-chain oxysterols, synthesized through the enzymatic or non-enzymatic oxidation of cholesterol, are appealing alternatives to cholesterol because of their lower abundance, higher hydrophilicity and structural similarity to cholesterol (Corcoran and Scott 2006; Dwyer et al. 2007). To address the issue of the endogenous lipidic activator of SMO, we took an unbiased genetic approach to identify lipid-related genes whose loss influences the strength of HH signaling.

## Results

### A focused CRISPR screen targeting lipid-related genes

Using our previously described strategy (Pusapati, Kong, Patel, Krishnan, et al. 2018) to identify positive and negative regulators of the HH pathway, we conducted focused loss-of-function CRISPR screens using a custom library targeting 1,244 lipid-related genes compiled by the LIPID MAPS consortium (**Supplementary Table 1** provides a list of all genes and guide RNAs in the library). This CRISPR library targeted all annotated genes encoding enzymes involved in the synthesis or metabolism of lipids as well as proteins that bind to or transport lipids. We used a previously characterized NIH/3T3 cell line (NIH/3T3-CG) that expresses <u>C</u>as9 and <u>G</u>FP driven by a HH-responsive fluorescent reporter (GLI-GFP) (Pusapati, Kong, Patel, Krishnan, et al. 2018). To ensure that HH signaling in these cells would be sensitive to perturbations of endogenous lipid metabolic pathways, the entire population of mutagenized cells was grown in lipoprotein-depleted media for one week prior to the screen and then treated with U18666A, a drug that prevents absorption of exogenous cholesterol from the lysosome, to minimize any sterol uptake from the media (**Fig.1A**). In the screen for positive regulators (hereafter the “HiSHH-Bot10%” screen), we treated cells with a saturating concentration of the ligand <u>S</u>onic <u>H</u>edge<u>h</u>og (HiSHH) and used Fluorescence Activated Cell Sorting (FACS) to collect poor responders, those with the lowest 10% of GLI-GFP fluorescence (**Figs.1A and 1B**; full screen results in **Supplementary Table 2**). In the screen for negative regulators, we treated cells with a low, sub-saturating concentration of SHH (LoSHH) that activated the reporter to <10% of maximal strength and selected super-responders, cells with the top 5% of GLI-GFP fluorescence (“LoSHH-Top5%” screen) (**Figs.1A and 1C;** full screen results in **Supplementary Table 3**).

**Figure 1.**
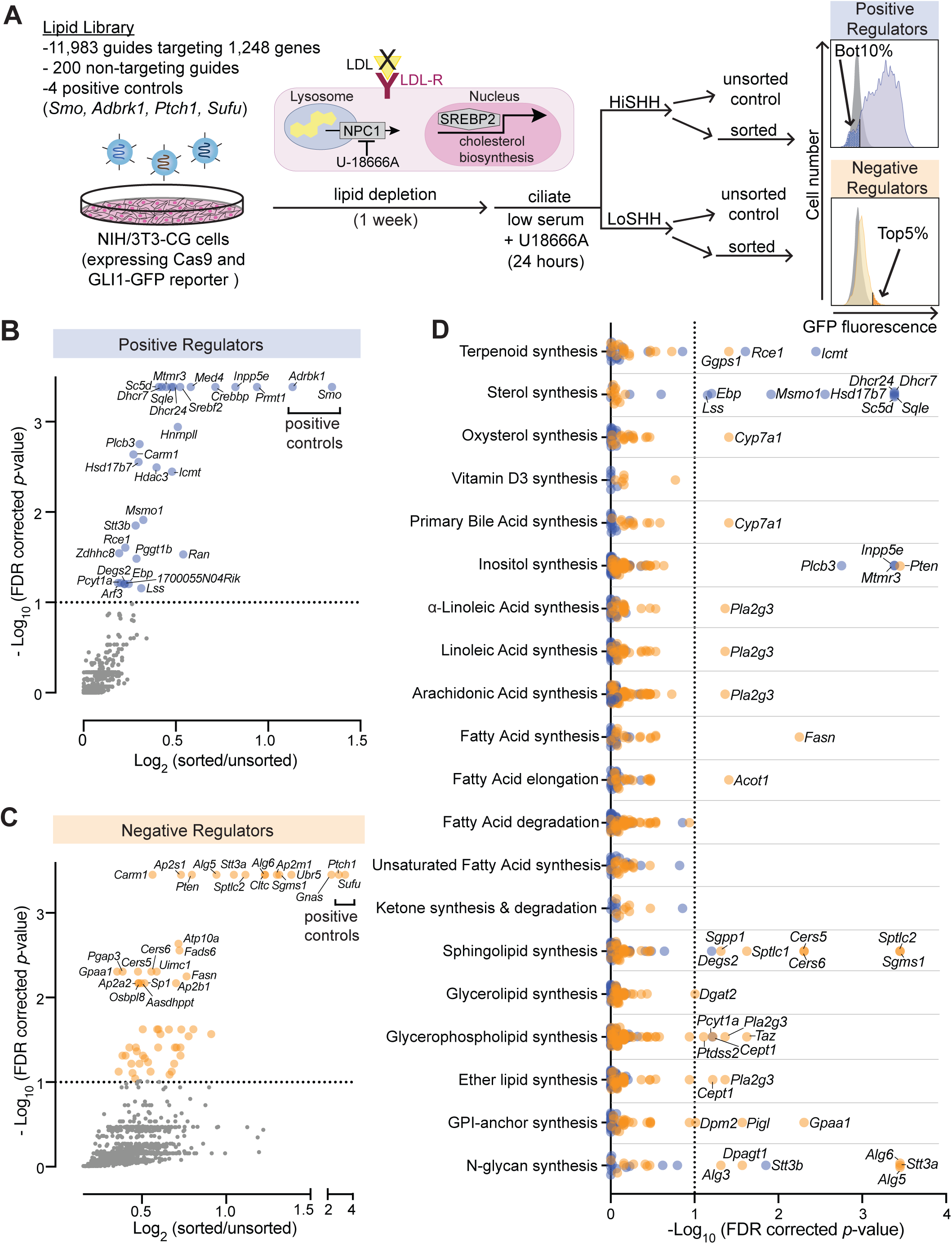
CRISPR screens identify lipid-related genes that influence Hedgehog signaling. (A) Flowchart summarizing the screening strategy. Inset highlights the dual strategy of lipid depletion and lysosomal cholesterol import inhibition used during the screen. (B and C) Volcano plots of the HiSHH-Bot10% (B) screen for positive regulators and the LoSHH-Top5% (C) screen for negative regulators. Enrichment is calculated as the mean of all sgRNAs for a given gene in the sorted over unsorted population, with the y-axis showing significance based on the false discovery rate (FDR)-corrected *p*-value. (D) Screen results analyzed by grouping genes based on the core lipid biosynthetic pathways in KEGG. In all panels, genes identified as positive and negative regulators are labeled in blue and orange respectively.

The screens correctly identified all four positive controls included in the library: *Smo* and *Adrbk1* (or *Grk2*) as positive regulators and *Ptch1* and *Sufu* as negative regulators (**Figs.1B and 1C**). In addition, genes previously known to influence HH signaling (*Gnas*) and protein trafficking at primary cilia (*Inpp5e*) were amongst the most significant hits (Regard et al. 2013; Garcia-Gonzalo et al. 2015; Chávez et al. 2015). In addition to *Inpp5e*, other genes involved in phosphoinositide metabolism (*Mtmr3* and *Plcb3*) were also significant hits. *Pla2g3*, which encodes a secreted phospholipase, was identified as a negative regulator of HH signaling, an effect that may be related to its known role as a suppressor of ciliogenesis (**Fig.1D**)(Gijs et al. 2015; Kim et al. 2010).

To identify lipid species that influence HH signaling, we separately analyzed the intersection of all genes expressed in NIH/3T3-CG cells based on RNAseq and annotated as part of a lipid metabolic pathway in the Kyoto Encyclopedia of Genes and genomes (KEGG) (**Fig.1D**; gene lists used for each pathway are shown in **Supplementary Table 4**; RNAseq data in **Supplementary Table 5**). Statistically significant hits clustered in two major pathways: (1) genes encoding enzymes in the cholesterol biosynthesis pathway were positive regulators of HH signaling and (2) genes encoding enzymes in the sphingolipid biosynthesis pathway were negative regulators (positive regulators are shown in blue and negative regulators in orange in **Fig.1D**). We focused on these two pathways for the work described in the rest of this study.

### Enzymes in the cholesterol biosynthesis pathway positively regulate Hedgehog signaling

Mutations in *Dhcr7* and *Sc5d*, which encode enzymes that catalyze the terminal steps in cholesterol biosynthesis, impair HH signaling in target cells and cause the congenital malformation syndromes Smith-Lemli-Opitz and lathosterolosis, respectively (Cooper et al. 2003; Blassberg et al. 2016; Porter and Herman 2011; Horvat, McWhir, and Rozman 2011). In addition to these genes, most of the genes encoding enzymes in the pathway that converts squalene to cholesterol were statistically significant hits with a FDR-corrected *p*-value threshold of 0.1 in the HiSHH-Bot10% screen (**Fig.2A**). Of the post-squalene cholesterol biosynthesis genes that did not meet this threshold, *Nsdhl* came close (FDR-corrected *p*-value = 0.25, **Supplementary Table 2**) and *Tm7sf2* is redundant (L. J. Sharpe and Brown 2013), leaving *Cyp51* as the only gene that was not identified. CRISPR-mediated loss-of-function mutations in *Lss*, required for an early step in the pathway, and in *Dhcr7* and *Dhcr24*, required for the terminal steps, impaired the transcriptional induction of endogenous *Gli1* (an immediate target gene used as a measure of signaling strength) (**Figs.2B and 2C**). HH signaling in both *Lss^-/-^* and *Dhcr7^-/-^* cells, but not in *Dhcr24^-/-^* cells, could be rescued with the addition of exogenous cholesterol, pointing to cholesterol deficiency as the cause of impaired HH signaling (**Figs.2B and 2C**). Rescue of HH signaling defects in *Dhcr7^-/-^* cells by exogenous cholesterol has also been demonstrated previously (Blassberg et al. 2016). We do not yet understand the inability of cholesterol to rescue signaling in *Dhcr24^-/-^* cells, but plan to address this unexpected observation with a dedicated study in the future. Nevertheless, since DHCR24 is only used in the Kandutsch-Russell and Bloch pathways for cholesterol synthesis but not in the shunt pathway (**Fig. 2A**), the defect in HH signaling in *Dhcr24^-/-^* cells suggests that products of the shunt pathway are not major regulators of HH signaling (L. J. Sharpe and Brown 2013).

**Figure 2.**
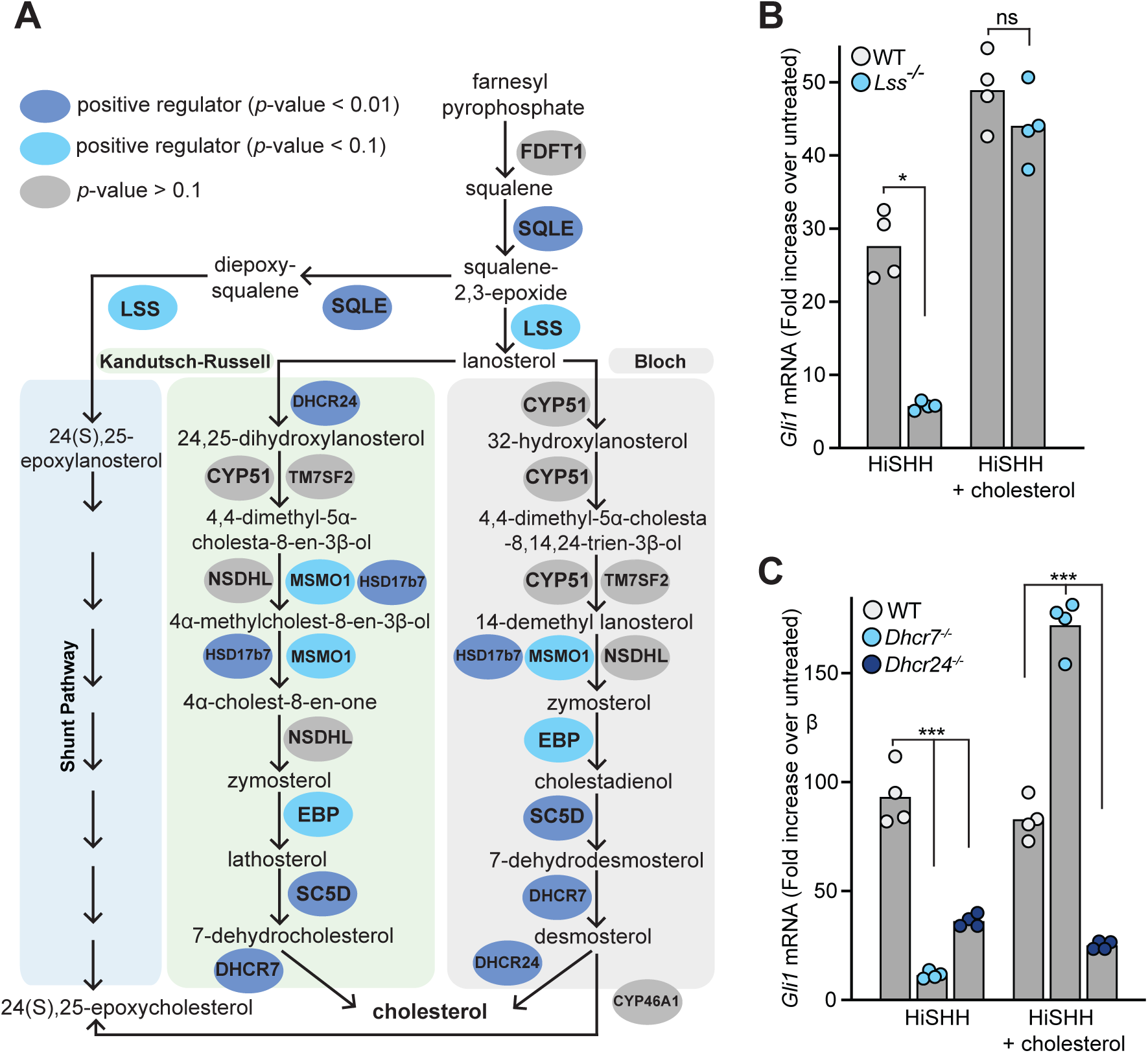
Enzymes that generate cholesterol positively regulate Hedgehog signaling. (A) The post-squalene portion of the cholesterol biosynthetic pathway, with enzymes colored according to their FDR corrected *p*-value in our CRISPR screens (see Supplementary Tables 2 and 3). Two branches of the pathway (the Kandutsch-Russel and the Bloch pathway) produce cholesterol, while a third shunt pathway does not require DHCR24 and produces 24(S),25-epoxycholesterol. (B and C) HH signaling strength in *Lss*^-/-^, *Dhcr7*^-/-^ and *Dhcr24*^-/-^ NIH/3T3 cells was assessed by measuring *Gli1* mRNA by quantitative reverse transcription PCR (qRT-PCR) after treatment with either HiSHH (25 nM) or HiSHH combined with 0.3 mM cholesterol: methyl-β-cyclodextrin (MβCD) complexes (see Methods for cholesterol: MβCD complexation). Statistical significance was determined by the Kruskal-Wallis test (B) or the Mann-Whitney test (C); \**p*-value≤ 0.05, \*\*\**p*-value≤ 0.001, non-significant (ns) *p*-value> 0.05.

The results of our unbiased screen highlight the importance of the endogenous post-squalene cholesterol biosynthetic pathway for HH signaling in target cells. While a simple explanation for this requirement is that cholesterol activates SMO in response to HH ligands, two additional possibilities have been discussed in the literature. First, defects in the terminal steps in cholesterol biosynthesis may lead to accumulation of precursor sterols that inhibit signaling (Porter and Herman 2011). However, HH signaling defects caused by mutations in genes that control the earliest steps in the pathway (*Sqle*, *Lss*; **Fig.2A**) cannot be explained by the accumulation of inhibitory precursor sterols (at least not by intermediates in the synthesis of cholesterol from squalene). The second possibility is that cholesterol is not the product of this pathway directly relevant to HH signaling. Instead, a different molecule synthesized from cholesterol, such as an oxysterol, primary bile acid or Vitamin D derivative, is required (Corcoran and Scott 2006; Dwyer et al. 2007; Bijlsma et al. 2006). However, none of the genes encoding enzymes that mediate synthesis of these metabolites (listed in **Supplementary Table 4**) were identified as significant hits in the HiSHH-Bot10% screen (**Fig.1D**). A lone oxysterol synthesis enzyme (CYP7A1) was implicated in an opposite role, a negative regulator, in the LoSHH-Top5% screen (**Fig.1D**).

In summary, the data from our genetic screen supports the view that cholesterol itself, rather than a precursor or a metabolite, is the endogenous sterol lipid that regulates SMO activation. Caveats of genetic screens include their inability to identify genes or pathways that are (1) redundant, (2) required for cell viability or growth, or (3) dependent on non-enzymatic reactions or exogenous molecules supplied by the media.

### Cellular sphingomyelin suppresses Hedgehog signaling

Multiple enzymes in the sphingolipid synthesis pathway were statistically significant hits in the LoSHH-Top5% screen, indicating these enzymes are negative regulators of HH signaling strength (**Fig.3A**). Top hits from this screen include *Sptlc2*, the first committed step in sphingolipid synthesis from serine and palmitoyl-CoA, as well as *Sgms1*, which converts ceramide to <u>s</u>phingo<u>m</u>yelin (SM). The identification of *Sgms1* suggests that SM is the relevant product of the sphingolipid pathway that attenuates HH signaling.

**Figure 3.**
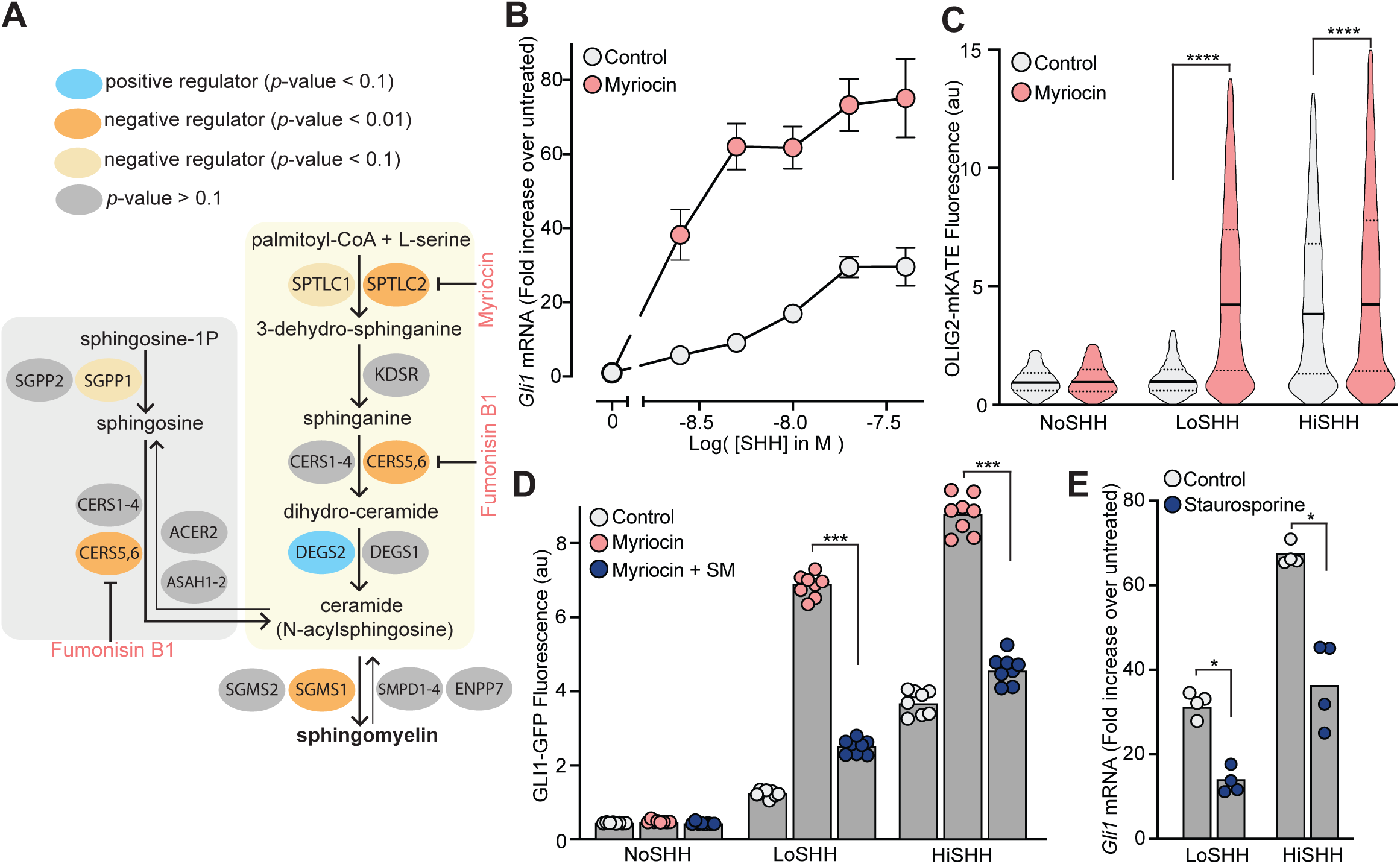
Enzymes that generate sphingomyelin negatively regulate Hedgehog signaling. (A) The pathway for the synthesis of SM, with enzymes colored according to their FDR corrected *p*-value in our CRISPR screens. (B) A dose-response curve for SHH in myriocin-treated NIH/3T3 cells compared to control cells treated with vehicle (DMSO) alone. Error bars denote standard deviation (n=4). (C) Differentiation of spinal Neural Progenitor Cells (NPCs) into OLIG2-postive motor neuron progenitors exposed to either LoSHH (5 nM) or HiSHH (25 nM) was assessed using flow cytometry to measure the fluorescence of a OLIG2-mKate differentiation reporter (n > 5000 cells for each treatment). (D) HH signaling strength in NIH/3T3-CG Reporter cells treated with LoSHH (5 nM) or HiSHH (50 nM) after treatment with myriocin alone or myriocin followed by addition of exogenous egg SM. Each data point represents the mean GLI1-GFP fluorescence derived from ∼250 cells. (E) HH signaling strength measured in NIH/3T3 cells treated with either LoSHH (5 nM) or HiSHH (25 nM) in the presence or absence of 50 nM staurosporine to increase SM. Statistical significance was determined by the Mann-Whitney test (C, D and E); *p-value≤ 0.05, ***p-value≤ 0.001, ****p-value≤ 0.0001.

Because we were unable to isolate viable NIH/3T3 cell lines entirely depleted of SPTLC2 or SGMS1 protein using CRISPR editing, we used an established pharmacological strategy. Myriocin is a fungal antibiotic that potently inhibits SPTLC2 (**Fig.3A**) and is commonly used to deplete SM in cells (Courtney et al. 2018; Tafesse et al. 2013, 2015). SM depletion by myriocin in NIH/3T3 cells was confirmed using both thin-layer chromatography (**Supplementary Fig.3A**) and flow cytometry of intact cells stained with a fluorescent protein probe (OlyA_E69A) that binds to total SM on the outer leaflet of the plasma membrane (**Supplementary Fig.3B**) (Endapally et al. 2019). Myriocin treatment markedly potentiated the response to SHH in NIH/3T3 cells, as measured by the transcriptional induction of *Gli1* (**Fig.3B**). This effect was also observed in two additional cell types. In mouse embryonic fibroblasts (MEFs), myriocin was sufficient to activate HH signaling even in the absence of added HH ligands (**Supplementary Fig.3C**). Mouse spinal neural progenitor cells (NPCs) differentiate into *Olig2*-expressing motor neuron progenitors in response to moderate concentrations of SHH. Myriocin potentiated the effect of SHH on NPCs, substantially reducing the concentration of SHH required to drive motor neuron differentiation (**Fig.3C**).

Several control experiments established that the potentiating effect of myriocin on HH signaling was caused by the depletion of SM, rather than an unrelated effect. Fumonisin B1, a mycotoxin structurally distinct from myriocin that inhibits a different step in SM synthesis (**Fig.3A**), also amplified HH signaling (**Supplementary Fig.3D**). Second, the potentiating effect of myriocin on HH signaling could be reversed by the exogenous administration of SM (**Fig.3D**). Finally, increasing SM levels in cells using low-dose staurosporine had the predicted opposite effect: reduction of HH signaling strength (**Fig.3E** and **Supplementary Fig.3E**) (Maekawa et al. 2016).

### Sphingomyelin restrains Hedgehog signaling at the level of Smoothened

SM is mostly localized in the outer leaflet of the plasma membrane where it plays key roles in its lateral organization, including the formation of ordered membrane microdomains that can influence protein trafficking, signaling and other processes (Simons and Ikonen 2000, 1997). Therefore, we looked broadly at the effects of myriocin on HH-relevant phenotypes, paying particular attention to primary cilia, organelles that are required for HH signaling in vertebrates (Huangfu et al. 2003). Myriocin did not significantly alter the abundances of the HH pathway proteins GLI3, SMO, SUFU or PTCH1 (**Supplementary Fig.4A**) and also did not change either the frequency or the length of primary cilia (**Supplementary Fig.4B**). Sensitivity of target cells to HH ligands can be influenced by the ciliary abundances of PTCH1, the receptor for all HH ligands that inhibits SMO, and by GPR161, a G-protein coupled receptor (GPCR) known to negatively regulate HH signaling (Rohatgi, Milenkovic, and Scott 2007; Pusapati, Kong, Patel, Gouti, et al. 2018; Mukhopadhyay et al. 2013). However, both proteins were properly localized in the ciliary membrane in myriocin-treated cells and were cleared (as expected) from cilia in response to SHH addition (**Supplementary Figs.4C and 4D**). Thus, myriocin does not seem to significantly alter ciliary biogenesis, ciliary morphology or ciliary trafficking events of receptors that negatively regulate HH signaling.

**Figure 4.**
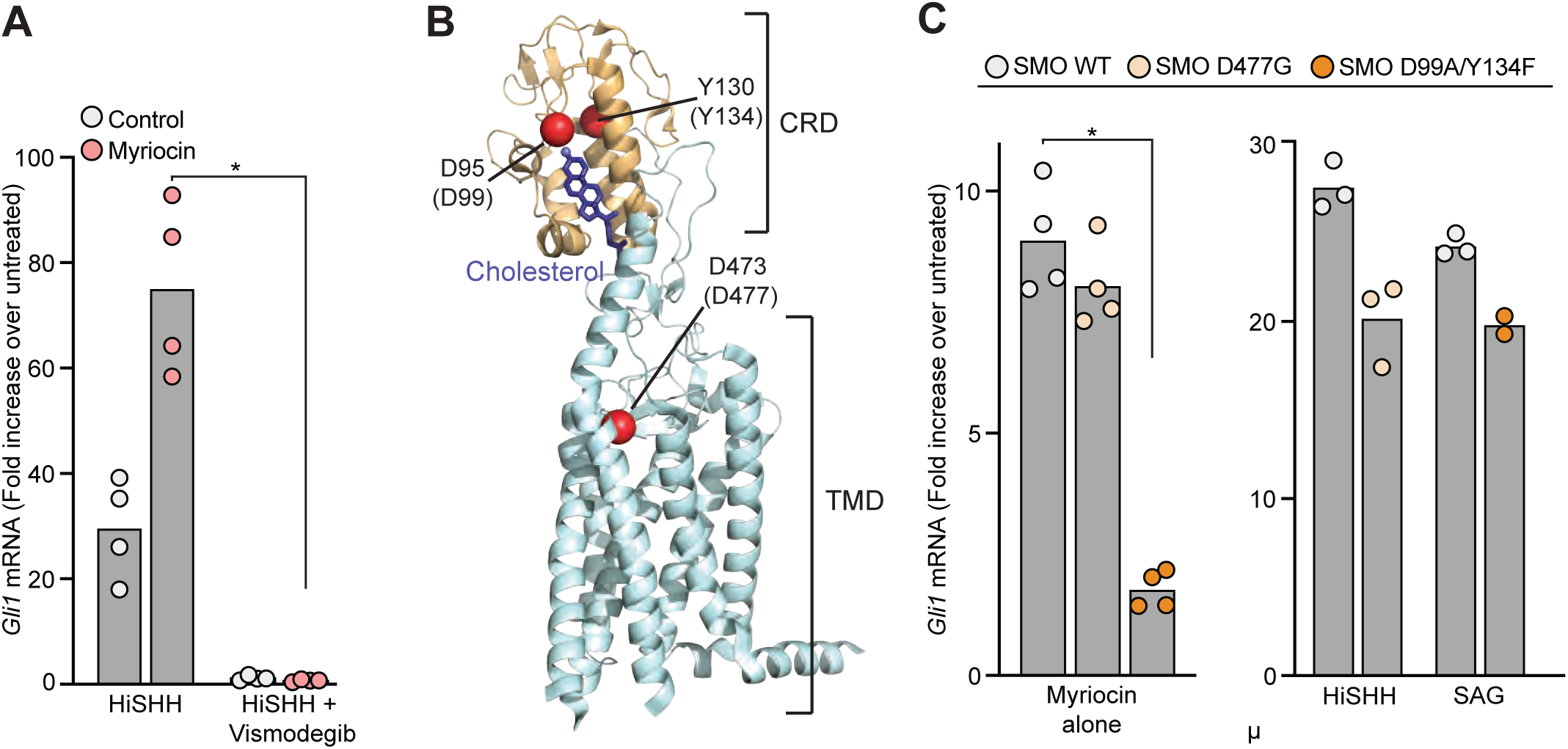
Sphingomyelin depletion potentiates Hedgehog signaling at the level of Smoothened. (A) HH signaling triggered by HiSHH (25 nM) in the presence or absence of Vismodegib (2.5 uM) in NIH/3T3 cells treated with myriocin. (B) Human SMO in complex with cholesterol (PDB 5L7D) highlighting two residues in the CRD (D95 and Y130) critical for cholesterol binding and a residue (D473) in the transmembrane domain (TMD) critical for binding to TMD ligands such as SAG. Numbering for mouse SMO, used in our studies, is denoted in parenthesis. (C) HH signaling triggered by myriocin alone in *Smo*^-/-^ MEFs stably expressing the indicated variants of mouse SMO. A control experiment (right) shows that the SMO variants respond appropriately to either SAG (100 nM) or HiSHH (25 nM), demonstrating protein integrity. Note that the D477G and D99A/Y134F mutations abrogate responses to SAG and SHH, respectively. Statistical significance was determined by the Mann-Whitney test (A and C); \**p*-value≤ 0.05.

The SHH-triggered accumulation of SMO in primary cilia is required for initiation of HH signaling in the cytoplasm (Corbit et al. 2005). Myriocin potentiated SMO ciliary accumulation in NIH/3T3 cells, suggesting that SM depletion enhances SMO activation (**Supplementary Fig.4E**). HH signaling in cells treated with myriocin was blocked by the SMO antagonist Vismodegib (**Fig.4A**), consistent with an effect on SMO or at a step upstream of SMO. The lack of an effect of myriocin on PTCH1 trafficking (**Supplementary Fig.4C**) led us to focus on SMO as the target for HH potentiation.

SMO has two well-defined ligand binding sites: one in the extracellular <u>c</u>ysteine-rich <u>d</u>omain (CRD) that binds cholesterol and oxysterols, and one in the trans<u>m</u>embrane <u>d</u>omain (TMD) that binds to diverse SMO agonists and antagonists (Eamon Fx Byrne et al. 2018; H. J. Sharpe et al. 2015). Another sterol binding site has also been recently described deep in the TMD (Deshpande et al. 2019). Mutations in the CRD site abolish sterol binding and responses to SHH; mutations in the more superficial TMD site prevent SMO activation by synthetic agonists like SAG but do not affect responses to SHH (**Fig.4B**) (E. F. Byrne et al. 2016; Luchetti et al. 2016; P. Huang et al. 2016). We tested the effects of these mutations on the ability of myriocin to activate HH signaling in *Smo^-/-^* MEFs stably expressing SMO variants. Since HH signaling is activated in these cells in response to myriocin alone (**Supplementary Fig.3C**), we were able to assess the effects of these mutations without the confounding effects of HH ligands or SMO agonists. Previously defined mutations in the sterol-binding CRD site (D99A/Y134F, **Fig.4B**), which abrogate cholesterol binding by disrupting a key hydrogen bond with the 3β-hydroxyl of cholesterol, prevented myriocin from activating signaling (**Fig.4C**). In contrast, a mutation (D477G, **Fig.4B**) in the TMD site failed to diminish myriocin-induced signaling (**Fig.4C**). In control experiments, SMO-D99A/Y134F and SMO-D477G were responsive to SAG and SHH, respectively, demonstrating protein integrity (**Fig.4C**). The fact that point mutations in SMO abrogated the effect of SM depletion suggests that myriocin influences HH signaling at the level of SMO.

### Sphingomyelin restrains Hedgehog signaling by sequestering cholesterol

The observation that mutations in the CRD of SMO, a well-defined binding site for cholesterol (E. F. Byrne et al. 2016), abrogated the effects of SM depletion (**Fig.4C**) suggested that SM regulates SMO activity by controlling the availability of cellular cholesterol that can bind to SMO. There is a precedent for SM regulation of cholesterol availability in another cellular signaling context, namely the control of cholesterol synthesis (Slotte and Bierman 1988; Scheek, Brown, and Goldstein 1997; Akash Das et al. 2014). These studies have led to a proposal that plasma membrane cholesterol is organized in three pools: a fixed pool essential for membrane integrity, a SM-sequestered pool with low chemical activity and a third (“accessible”) pool with higher chemical activity that is available to interact with proteins (Akash Das et al. 2014; A. Radhakrishnan, Anderson, and McConnell 2000; Y. Lange, Cutler, and Steck 1980; Yvonne Lange, Ye, and Steck 2004). The distribution of cholesterol between the sequestered and accessible pools is determined by the ratio of cholesterol to SM: SM depletion (or cholesterol addition) lead to an increase in accessible cholesterol (**Fig.5A**) (Akash Das et al. 2014). These different pools of cholesterol are characterized by differences in the chemical activity (or accessibility) of membrane cholesterol and thus cannot be measured by mass spectrometry, which requires the extraction of total cholesterol from cell membranes with solvents (Steck and Lange 2010; McConnell and Radhakrishnan 2003). Instead, protein probes derived from pore-forming toxins have been recently developed to detect the accessible and sequestered pools in intact membranes or cells (**Fig.5A**): PFO*, derived from the bacterial toxin <u>P</u>erfringolysin <u>O</u>, binds to the accessible pool of cholesterol and OlyA, derived from the fungal toxin <u>O</u>stero<u>ly</u>sin <u>A</u>, binds to SM-cholesterol complexes (Endapally et al. 2019; A. Das et al. 2013; Skočaj et al. 2014; Hotze et al. 2002). A useful point mutant of OlyA (OlyA_E69A) binds to both free SM and SM-cholesterol complexes, allowing measurement of total SM (**Fig.5A**) (Endapally et al. 2019). To test these probes in NIH/3T3 cells, we used flow cytometry to measure the binding of fluorescently-labeled PFO*, OlyA and OlyA_E69A to intact cells either treated with myriocin to deplete SM or loaded with exogenous cholesterol (known to increase accessible cholesterol levels in cells (Akash Das et al. 2014)). Treatment of cells with myriocin decreased both OlyA and OlyA_E69A staining, consistent with SM depletion (**Fig.5B and 5C**); staining was restored by the addition of exogenous egg SM (**Fig.5D**). Myriocin treatment and cholesterol loading increased PFO* staining, showing that both treatments increased the levels of accessible cholesterol in the plasma membrane (**Fig.5E**). The depletion of SM with myriocin or other pharmacological agents does not change levels of total cellular cholesterol or plasma membrane cholesterol (Tafesse et al. 2013, 2015; Akash Das et al. 2014).

**Figure 5.**
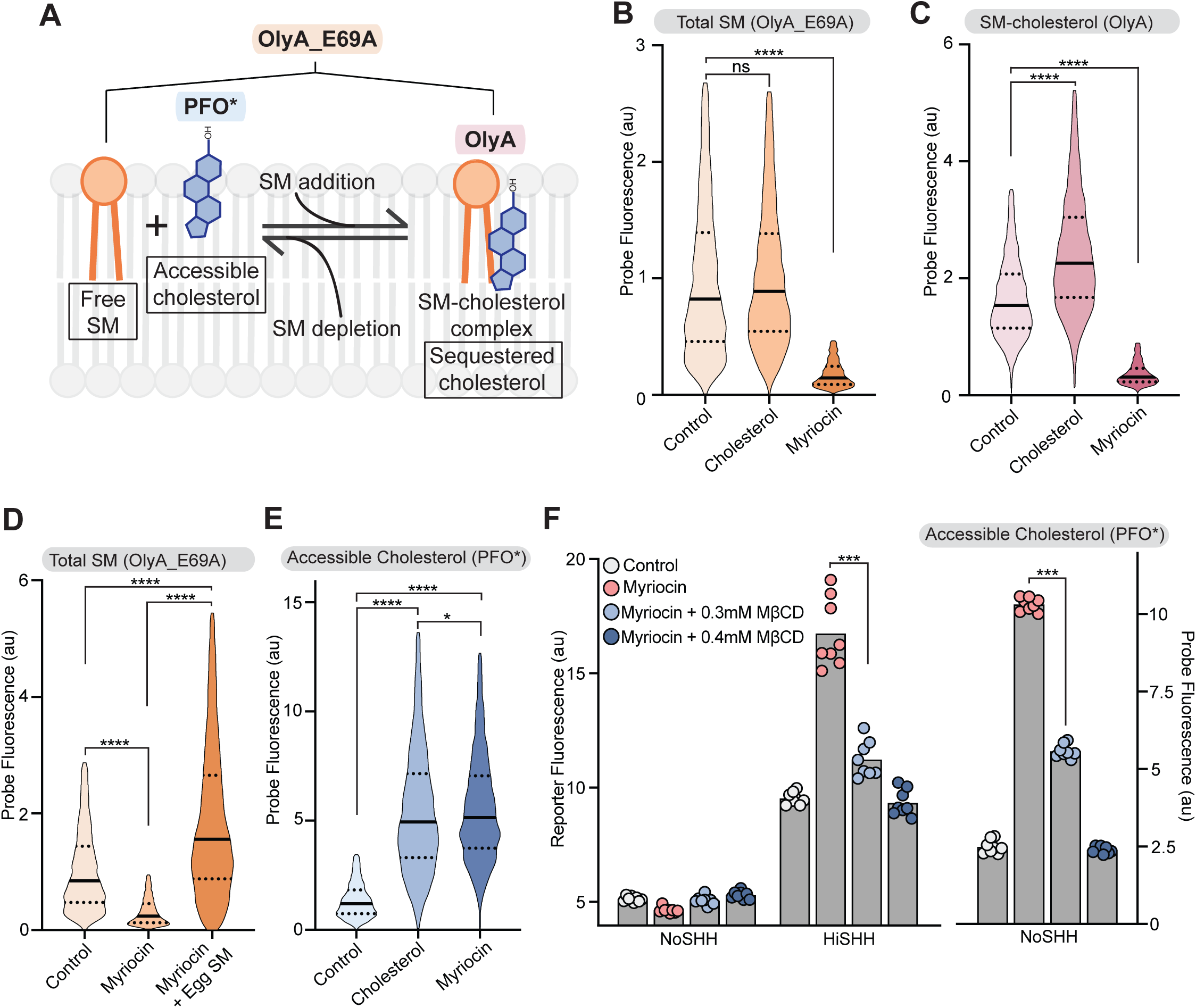
Reducing accessible cholesterol in myriocin-treated cells impairs Hedgehog signaling. (A) Cholesterol and SM form SM-cholesterol complexes in which cholesterol is sequestered and prevented from interacting with proteins like SMO. The ratio of SM to cholesterol determines the level of accessible cholesterol (free from SM). Protein probes detecting the various pools of cholesterol are shown: PFO* binds to accessible cholesterol, OlyA to SM-cholesterol complexes and OlyA_E69A to both free SM and SM-cholesterol complexes. (B-E) Flow cytometry of intact cells stained with fluorescently-labeled OlyA_E69A (B and D), OlyA (C) or PFO* (E) after the indicated treatments (n > 4000 cells for each condition). (F) HiSHH-induced (50 nM) GLI1-GFP reporter fluorescence in NIH/3T3-CG cells treated with myriocin alone or myriocin followed by two concentrations of MβCD to reduce accessible cholesterol levels. The graph on the right shows whole cell fluorescence of the same cells stained with PFO* to measure accessible cholesterol in the outer leaflet of the plasma membrane. Each data point denotes the mean fluorescence of GLI1-GFP or PFO* staining calculated from >250 cells. Statistical significance was determined by the Mann-Whitney test (B-F); \**p*-value≤ 0.05, \*\*\**p*-value≤ 0.001, \*\*\*\**p*-value≤ 0.0001, non-significant (ns) *p*-value> 0.05.

With these tools in hand, we sought to test the model that SM depletion by myriocin potentiates HH signaling by increasing the pool of accessible cholesterol. Reducing accessible cholesterol in myriocin-treated cells with methyl-β-cyclodextrin (MβCD), measured by PFO* staining, decreased SHH-induced activation of the GLI-GFP reporter back to levels seen in untreated cells (**Fig.5F**). This rescue is not consistent with the alternative possibility that SM negatively regulates SMO either directly or through a different mechanism. These results, together with the requirement of the cholesterol-binding CRD for the potentiating effect of myriocin (**Fig.4C**), support the model that SM impairs HH signaling by sequestering cholesterol into complexes where it is inaccessible to SMO. This conclusion is also consistent with the observation that purified SMO is constitutively active in nano-discs containing physiological levels of cholesterol in the absence of SM (Myers et al. 2017). In summary, the effects of SM depletion with myriocin are reminiscent of those when cells are loaded with cholesterol: accessible cholesterol levels increase and HH signaling is amplified (Akash Das et al. 2014; Luchetti et al. 2016; P. Huang et al. 2016).

### The ciliary membrane is a compartment with low cholesterol accessibility

The regulation of SMO by PTCH1 occurs at primary cilia, the only post-Golgi compartment in the cell where both proteins can be found localized together (Rohatgi et al. 2009; Rohatgi, Milenkovic, and Scott 2007). Since the ciliary membrane is thought to have a different lipid (and protein) composition than the plasma membrane (recently reviewed by (Nachury and Mick 2019)), we compared PFO*, OlyA and OlyA_E69A staining in the ciliary membrane relative to the plasma membrane using confocal microscopy. Controls confirmed that these probes could be used to measure levels of cholesterol, SM-cholesterol complexes and total SM in the ciliary membrane using quantitative fluorescence microscopy (**Supplementary Figs.6A-6C**), analogous to how we used them to measure these species at the plasma membrane by flow cytometry (**Figs.5B-5E**).

**Figure 6.**
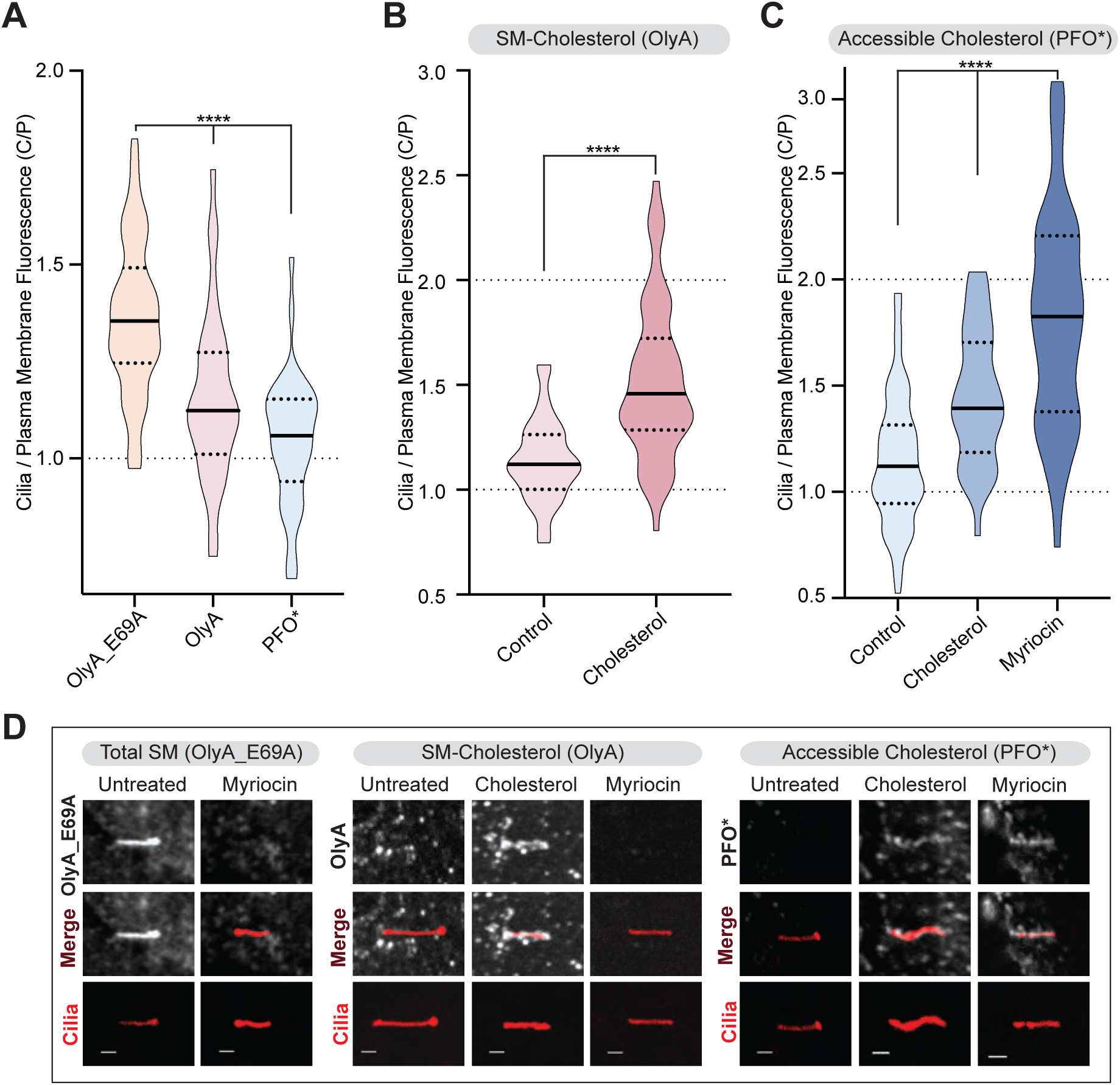
Primary cilia have high sphingomyelin and low accessible cholesterol. (A-C) Ratio of mean ciliary staining intensity to mean plasma membrane staining intensity (the C/P ratio, see text) for OlyA_E69A, OlyA, or PFO* (see Fig. 5A) in NIH/3T3 cells left untreated (A) or treated with either myriocin or cholesterol-MβCD complexes (B and C) (A-C, n > 30 cilia per condition). (D) Representative images of individual primary cilia from cells stained with each of the lipid probes after the indicated treatments. Cells stably expressed ARL13B-GFP to allow the identification of cilia. Scale bar: 1 micron. Statistical significance was determined by the Kruskal-Wallis test (A and C) or the Mann-Whitney test (B); \*\*\*\**p*-value≤ 0.0001.

For each cilium imaged, we calculated the ratio of mean ciliary fluorescence to mean plasma membrane fluorescence in a region surrounding the cilium (hereafter the “C/P ratio”)(Geneva, Tan, and Calvert 2017). We used this metric because myriocin treatment or cholesterol loading will lead to changes in probe staining at *both* the plasma membrane and the ciliary membrane. The C/P ratio reflects changes in the ciliary membrane *relative* to changes in the plasma membrane: if probe staining increases by the same factor in the plasma membrane and the ciliary membrane the C/P ratio will remain unchanged.

In untreated cells, the C/P ratio was significantly higher for OlyA_E69A staining compared to OlyA or PFO* staining (**Fig.6A**). This suggests that the ratio of SM to cholesterol, which determines the abundance of accessible cholesterol, is higher in the ciliary membrane compared to the plasma membrane. Indeed, while OlyA_E69A staining of total SM was readily detectable in cilia, most cilia did not show distinctive staining for SM-cholesterol complexes (OlyA) or accessible cholesterol (PFO*) (**Fig.6D**). Cholesterol loading increased the C/P ratio of SM-cholesterol complexes (OlyA staining, **Figs.6B and 6D**), showing that a significant proportion of SM molecules at cilia are free to pair with exogenously added cholesterol. Myriocin treatment, which reduced SM levels in both the plasma membrane and the ciliary membrane (**Fig.5B and Supplementary Fig.6A**), increased the amount of accessible cholesterol in the ciliary membrane relative to the plasma membrane (PFO* staining, **Fig.6C and 6D**). Indeed, myriocin had a greater effect on the C/P ratio of accessible cholesterol compared to even cholesterol loading, suggesting that SM provides the major restraint on cholesterol accessibility at cilia (**Fig.6C**). Taken together, we conclude that the ratio of sphingomyelin to cholesterol is higher in the ciliary membrane compared to the plasma membrane, leading to reduced cholesterol accessibility. This may be critical for keeping SMO, which is cycling through the ciliary membrane even in the absence of SHH, in an inactive state by restricting its access to cholesterol (Ocbina and Anderson 2008). Reducing SM levels with myriocin (or increasing cholesterol levels by cholesterol loading) increases accessible cholesterol in the ciliary membrane and, consequently, potentiates SMO activation.

### Hedgehog ligands cause an increase in cholesterol accessibility at primary cilia

PTCH1, the receptor for HH ligands, is thought to inhibit SMO by reducing its access to cholesterol using its transporter-like activity. Since PTCH1 is localized in and around the cilium, we have proposed that it could function to inhibit SMO by reducing accessible cholesterol in the ciliary membrane (Luchetti et al. 2016; Kong, Siebold, and Rohatgi 2019; P. Huang et al. 2016). This model predicts that SHH, which inhibits PTCH1 activity, should lead to an increase in accessible cholesterol and PFO* staining at the ciliary membrane. Given the potential artifacts associated with overexpressing a transporter protein, we sought to measure changes in accessible cholesterol at endogenous PTCH1 expression levels.

SHH did not induce much of a change in PFO* staining of the bulk plasma membrane, measured by FACS (**Fig.7A**). A lack of an effect is not surprising because changes in overall cholesterol accessibility would influence many other cellular processes, including the signaling system that maintains cholesterol homeostasis (Steck and Lange 2010; M. S. Brown and Goldstein 2009; Michael S. Brown, Radhakrishnan, and Goldstein 2018). In contrast, SHH led to a rapid increase in accessible cholesterol in the presence of myriocin in the ciliary membrane (**Figs.7B and 7C**). SHH did not change ciliary PFO* staining in *Ptch1^-/-^* cells (**Supplementary Fig.7A**). In addition, activation of signaling with SAG, which bypasses PTCH1 and directly activates SMO, also did not cause significant changes in accessible cholesterol at the ciliary membrane (**Supplementary Fig.7B**). Both controls show that the SHH-induced changes in accessible cholesterol at primary cilia are dependent on PTCH1 activity. In the absence of myriocin, we detected only minimal changes in PFO* staining in cilia (**Fig.7B**). We believe that this is related to a well-known property of PFO*: it only binds to membranes when cholesterol rises above a specific threshold and becomes accessible (A. Das et al. 2013; Flanagan et al. 2009; Sokolov and Radhakrishnan 2010). In untreated cells, accessible cholesterol levels at cilia are too low (even with SHH) to allow sufficient probe binding. However, SM depletion induced by myriocin raised accessible cholesterol levels above the threshold levels required for PFO* binding, thus allowing us to detect increases in ciliary accessible cholesterol induced by SHH.

**Figure 7.**
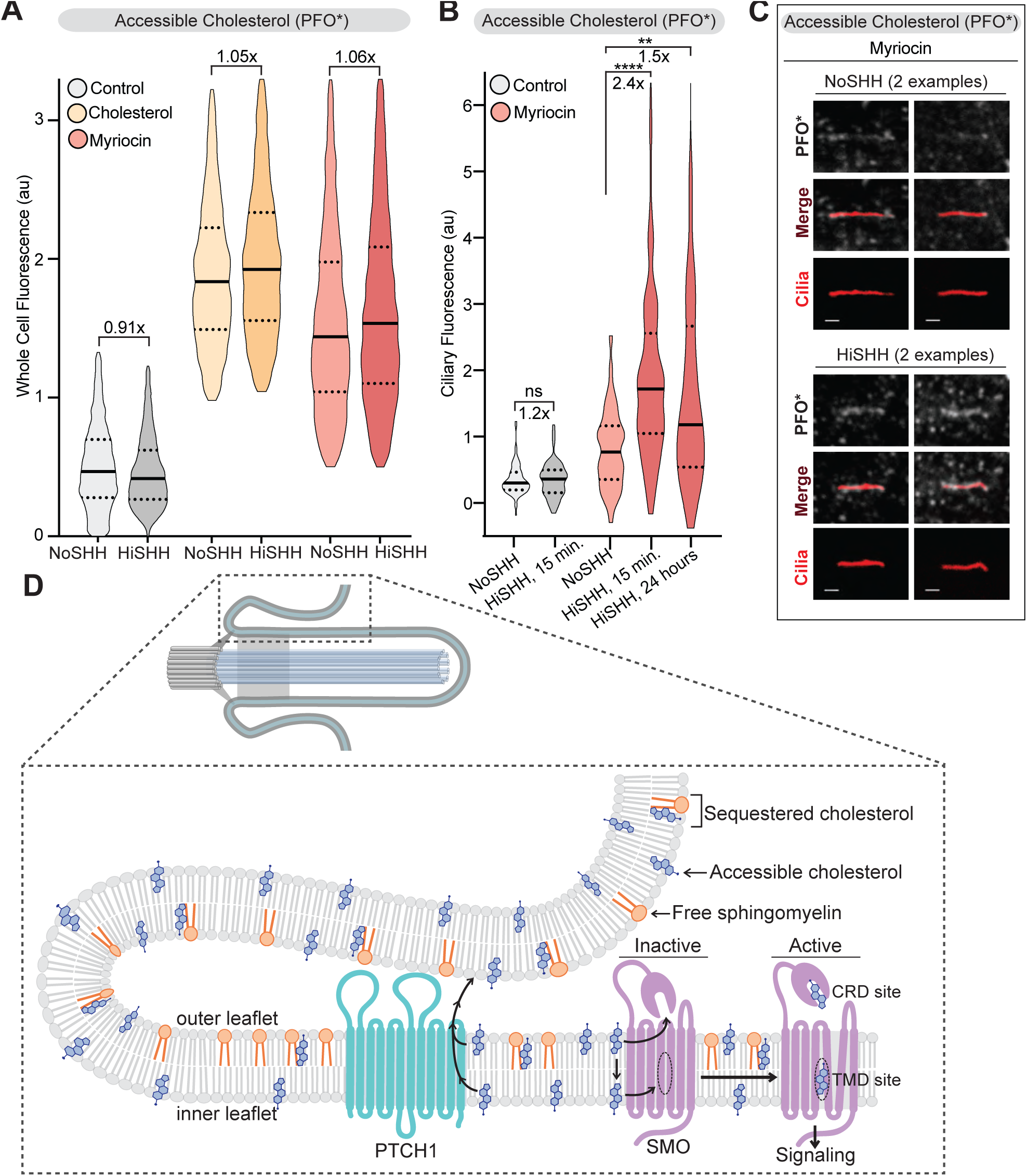
PTCH1 decreases the pool of accessible cholesterol at primary cilia. (A) Flow cytometry was used to measure plasma membrane PFO* staining in intact cells after HiSHH (25 nM) treatment (n > 4000 cells per condition). Fold changes of median values (SHH treated over untreated) are indicated (A and B). (B) PFO* staining at primary cilia in the presence of HiSHH (25 nM) in cells treated with myriocin or left untreated (n > 65 cilia per condition). (C) Representative images of primary cilia in cells treated with myriocin with or without the addition of HiSHH (25 nM). Scale bar: 1 micron. (D) Model depicting how PTCH1 could inhibit SMO at primary cilia by decreasing accessible cholesterol in the ciliary membrane (see Discussion for details). Statistical significance was determined by the Mann-Whitney test (B); \*\**p*-value≤0.01, \*\*\*\**p*-value≤0.0001, non-significant (ns) *p*-value>0.05.

## Discussion

The results of our unbiased screen for lipid-related genes that influence the strength of HH signaling uncovered two pathways-- cholesterol and SM synthesis-- that both converge on accessible cholesterol as the critical species that regulates the interaction between PTCH1 and SMO. The potentiating effect of SM depletion on HH signaling points to cholesterol itself as the regulatory sterol, since side-chain oxysterols do not form analogous complexes with SM (and the lipid probes used in our studies do not interact with oxysterols)(Endapally et al. 2019; Bielska et al. 2012; Gale et al. 2009). Two features explain how a seemingly abundant membrane lipid like cholesterol can play an instructive role as a second messenger in a signaling pathway. First, only a fraction of total plasma membrane cholesterol is relevant to the regulation of HH signaling. This thermodynamically distinct pool of accessible cholesterol with high chemical activity ranges from ∼2% of total plasma membrane cholesterol in lipid depleted cells to ∼15% of total plasma membrane cholesterol in lipid replete cells (Akash Das et al. 2014). Second, the changes in accessible cholesterol that regulate SMO are confined to a subcellular compartment, the primary cilium, where PTCH1, SMO and the cytoplasmic signaling machinery downstream of SMO are localized. This latter strategy of confining changes in abundance of a second messenger to a subcellular compartment or microdomain is commonly used in other signaling pathways, such as those that use cAMP or calcium. This feature of localization allows small changes in absolute numbers of a regulatory molecule to be translated into larger changes in concentration and also insulates other cellular processes from being inappropriately impacted (because second messengers are commonly used in multiple pathways).

The SCAP/SREBP signaling system that ensures tight control over cholesterol homeostasis provided an important precedent (and inspiration) for our work (Michael S. Brown, Radhakrishnan, and Goldstein 2018; M. S. Brown and Goldstein 2009). This system also senses accessible cholesterol in a specific subcellular compartment, the endoplasmic reticulum (ER), to regulate the transcription of genes that control cholesterol biosynthesis (Arun Radhakrishnan et al. 2008). Continual transport of cholesterol from the plasma membrane to the ER ensures that information about changes in plasma membrane cholesterol accessibility is transmitted to the ER, where the sensor proteins are located (Infante and Radhakrishnan 2017). Confining the effects of HH ligands on cholesterol accessibility to the ciliary membrane, rather than the plasma membrane, is required to prevent crosstalk with this important homeostatic pathway in the cell.

How might PTCH1 reduce the levels of accessible cholesterol in the ciliary membrane? In principle, PTCH1 could accomplish this either by increasing ciliary levels of SM or by decreasing ciliary levels of cholesterol. PTCH1 is more likely to be a sterol transporter for several reasons: it has homology to the cholesterol transporter NPC1 and the spate of recent PTCH1 structures have identified putative sterol ligands (Zhang et al. 2018; Gong et al. 2018; X. Qi et al. 2018; C. Qi et al. 2018; Qian et al. 2019). We have recently presented a model for how PTCH1 could reduce levels of cholesterol in the ciliary membrane by transporting it to the closely opposed membranes of the ciliary pocket (**Fig.7D**) (Kong, Siebold, and Rohatgi 2019). The level of SM in the ciliary membrane would determine the set-point for how much cholesterol needs to be transported by PTCH1 to reduce accessible cholesterol below the threshold required to activate SMO. Depleting SM (or loading cells with cholesterol) raises the demands on PTCH1 transport activity and hence makes cells hyper-sensitive to PTCH1-inactivating HH ligands. In some cell types (MEFs, see **Supplementary Fig.3C**), PTCH1 likely cannot overcome the effect of SM depletion, leading to SMO activation even in the absence of any SHH. The emphasis in this model on the SM-cholesterol ratio accounts for the observation that either SM depletion (**Fig.3B**) or cholesterol loading (Luchetti et al. 2016; P. Huang et al. 2016) synergizes with HH ligands to activate signaling.

SMO has two potential cholesterol-binding sites: one in the CRD (tested in **Fig.4C**) and a second deep in the TMD (E. F. Byrne et al. 2016; Luchetti et al. 2016; Pengxiang Huang et al. 2018; Deshpande et al. 2019). Mutations in either site prevent the activation of signaling by HH ligands, implicating both in the regulation of SMO by PTCH1. The extracellular CRD site is closer to the outer leaflet of the plasma membrane, while the TMD site has been proposed to obtain cholesterol from the inner leafter through an opening between transmembrane helices 5 and 6 (E. F. Byrne et al. 2016; Pengxiang Huang et al. 2018; Deshpande et al. 2019). Importantly, SM is confined to the outer leaflet of the plasma membrane and thus its depletion will directly increase accessibility of cholesterol in the outer leaflet (Steck and Lange 2018). Indeed, the probes used in our work are added to intact cells and hence monitor levels of accessible cholesterol, SM, and SM-cholesterol complexes in the outer leaflet. However, SM depletion likely also causes an increase in the accessibility of inner leaflet cholesterol because cholesterol can rapidly redistribute between the two leaflets of the plasma membrane by flip-flop movement (Steck and Lange 2018). Thus, SM depletion can increase cholesterol access to either the CRD or the TMD site in SMO (**Fig.7D**).

We end with a speculative answer to an enigma in HH signaling: why is the HH pathway dependent on primary cilia in vertebrates but not in *Drosophila*? The predominant sterol in *Drosophila* is ergosterol, with cholesterol itself representing <5% of membrane sterols (Rietveld et al. 1999). In addition, flies are cholesterol auxotrophs: they acquire cholesterol from the diet and have lost many of the genes for cholesterol biosynthesis (Vinci, Xia, and Veitia 2008; Carvalho et al. 2010). Thus, flies do not need the regulatory machinery that monitors accessible cholesterol in the plasma membrane and adjusts the transcription of cholesterol biosynthetic genes. Indeed, the SREBP pathway, which monitors accessible cholesterol in vertebrates, has been repurposed in *Drosophila* to respond to phosphatidylethanolamine (Dobrosotskaya et al. 2002). Cholesterol levels in *Drosophila* are instead sensed by a very different nuclear receptor-based mechanism (Bujold et al. 2010). We propose that the lack of a need to regulate cholesterol biosynthetic pathway genes abrogates the need to sequester HH signaling in primary cilia in insects.

## Methods

### Reagents

Suppliers for chemicals included Sigma-Aldrich (U18666A, cholesterol, Methyl-β-cyclodextrin, fatty acid free Bovine Serum Albumin (BSA), Atto-647N maleimide); Cayman Chemicals (Myriocin, Fumonisin B), Calbiochem (Staurosporine), Avanti Polar Lipids (Egg sphingomyelin), Matreya LLC (Milk sphingomyelin), Enzo Life Sciences (SAG), LC Labs (Vismodegib) and Invitrogen (Alexa-647 NHS ester). Primary (SMO, PTCH1, SUFU, GLI1, GLI3 and P38) and secondary antibodies (Peroxidase AffiniPure Donkey Anti-Mouse, Rabbit and Goat IgG (H+L)) used for Western Blotting (Supplementary Fig.4A) were previously described (Pusapati et al., 2018). Human SHH was expressed and purified as described previously (Bishop et al., 2009). Reagents used for cell culture are discussed in the methods for Cell Culture and Drug Treatments.

### Guide RNA library targeting lipid-related genes

In order to generate a lipid-library, human lipid-modifying genes/proteins were downloaded from the LIPID MAPS Proteome Database (“LIPID MAPS Proteome Database,” n.d.). Using Entrez IDs, these human gene names/IDs were converted to their mouse homologs using a database from the Mouse Genome Informatics (MGI). Any human genes not found in the MGI database were manually confirmed to not have a mouse homolog and excluded from the library, or, if a mouse homolog existed for the human gene, it was added to the target gene list. Missing genes were supplemented manually. This resulted in a list of 1,244 mouse lipid-related genes. Finally, *Ptch1*, *Sufu*, *Smo*, and *Adrbk1* (*Grk2*) were added to the target gene list as positive controls. Using lists from the Brie Mouse CRISPR Knockout Pooled library (Doench et al. 2016) and the GeCKO v2 Mouse CRISPR Knockout Pooled library (Sanjana, Shalem, and Zhang 2014), guide RNA (sgRNA) sequences were extracted. Genes for which guides could be found in either the Brie or GeCKO library, the guide count per gene ranged from 4 to 10. Approximately 15 genes had no guides in the Brie or GeCKO libraries. For these genes, 10 guide sequences were designed using the Broad Institute’s CRISPR Design tool (https://portals.broadinstitute.org/gpp/public/analysis-tools/sgrna-design). The final target gene list contained 11,783 guides targeting 1,248 genes. 200 non-targeting control guides were added to this final list from the GeCKO v2 library, resulting in a total of 11,983 guides for the library. The library of guides was synthesized using Twist Bioscience’s Oligo Pools and cloned into lentiGuide Puro vector (a gift from Feng Zhang; Addgene plasmid #52963) as described previously (Joung et al. 2017).

### CRISPR/Cas9 screen targeting lipid-related genes

Our reporter-based screening platform has been described previously in detail (Lebensohn et al. 2016; Pusapati, Kong, Patel, Krishnan, et al. 2018). NIH/3T3-CG cells were used because they respond to SHH in a concentration-dependent manner, carry stably integrated Cas9, and carry fluorescence-based, quantitative reporter of HH signaling (GLI-GFP) (Pusapati, Kong, Patel, Krishnan, et al. 2018). This reporter allows the isolation of cell populations with enhanced or reduced HH signaling phenotypes by FACS. CRISPR library amplification, lentiviral production, functional titer determination and transduction were carried out as previously described in detail (Joung et al. 2017; Pusapati, Kong, Patel, Krishnan, et al. 2018).

To prepare the library of cells for screening, 4x 15 cm plates were seeded with 5 million cells each. These cells were then grown for 1 week in “supplemented DMEM” (see methods section on Cell Culture and Drug Treatments) containing 5% Lipoprotein Depleted Serum (LDS) in replace of 10% Fetal Bovine Serum. This treatment was carried out so that cells would become reliant on their own endogenous lipid-biosynthesis machinery. Finally, cells were grown to confluence in 5% LDS DMEM and then serum starved in 0.5% LDS DMEM and treated with 1 μM U18666A and either left untreated (NoSHH) or treated with LoSHH (3.2 nM) or HiSHH (25 nM) for 24 hours. Cells were then trypsinized, 4 million were pelleted and frozen for the unsorted control population, and the remaining ∼25 million cells (representing 2000-fold coverage of the sgRNA library) were sorted for the lowest 10% (HiSHH-Bottom10% screen) or highest 5% of GFP fluorescence (HiSHH-Top5% screen). Finally, genomic DNA was extracted from the unsorted and sorted cells and the sgRNA library was amplified by nested PCR, subjected to Illumina sequencing, and analyzed using the MAGeCK algorithm as described previously (Pusapati, Kong, Patel, Krishnan, et al. 2018; Li et al. 2014).

### Kyoto Encyclopedia of Genes and genomes (KEGG) analysis of Lipid Pathways

In order to determine which lipids influence Hedgehog signaling, mouse-specific genes were manually curated into lists for each lipid metabolic pathway identified on the Kyoto Encyclopedia of Genes and genomes (KEGG) website (**Supplementary Table 4**). Since oxysterol pathways are not a separate category in KEGG, manual curation of the literature was used to identify 34 oxysterol-related enzymes (see **Supplementary Table 4**), 6 of which were not found in the KEGG database (Sever et al. 2016; Griffiths, Crick, et al. 2019; Griffiths, Yutuc, et al. 2019; Abdel-Khalik et al. 2018; Raleigh et al. 2018; Griffiths and Wang 2018; Griffiths et al. 2017). For each gene identified in KEGG or the oxysterol list, FDR-corrected p-values were extracted from the HiSHH-Bot10% screen (Supplementary Table 2) as well as the LoSHH-Top5% screen (Supplementary Table 3). Finally, the expression level of each gene was obtained from RNAseq analysis (performed in duplicate and RPKM normalized) carried out in NIH/3T3 cells (**Supplementary Table 5**). If a gene was not expressed (a value of 0 in both RNAseq data sets), it was not included in the pathway analysis of Fig1D.

### Cell Culture and Drug Treatments

NIH/3T3-CG Reporter Cells used in the CRISPR screen (and in **Figs. 3D** & **5F**), Smo-/- MEFs stably expressing SMO mutants (WT, D477G and D99A/Y134F), and NIH/3T3 Flp-In cells stably expressing GPR161-YFP have been previously described and characterized (Pusapati, Kong, Patel, Krishnan, et al. 2018; Luchetti et al. 2016; Pusapati, Kong, Patel, Gouti, et al. 2018). NIH/3T3 Flp-In cells were purchased from Thermo Fisher Scientific and NIH/3T3 cells from ATCC. In the NIH/3T3 Flp-In background, Lss-/-, Dhcr7-/- and Dhcr24-/- clonal cell lines were generated using a two-cut CRISPR strategy using methods described in our previous publications (Pusapati, Kong, Patel, Krishnan, et al. 2018; Pusapati, Kong, Patel, Gouti, et al. 2018). NIH/3T3 cells and *Ptch^-/-^* MEFs (Rohatgi, Milenkovic, and Scott 2007) stably expressing ARL13B C-terminally tagged with GFP were generated by lentiviral infection followed by puromycin (Calbiochem) selection.

All cells were grown in high glucose Dulbecco’s Modified Eagle’s Medium (DMEM) (Thermo Fisher Scientific) containing the following supplements (hereafter referred to as “supplemented DMEM”): 10% Fetal Bovine Serum (FBS) (Sigma), 1 mM sodium pyruvate (Gibco), 2 mM L-glutamine (Gemini Biosciences), 1× MEM nonessential amino acids solution (Gibco), penicillin (40 U/ml) and streptomycin (40 g/ml) (Gemini Biosciences). To induce ciliation, cells were grown to confluence and then the cell media was exchanged to low serum (0.5% FBS) supplemented DMEM. In order to test the requirement for cholesterol in Hedgehog signaling, *Dhcr7^-/-^* and *Dhcr24^-/-^* NIH/3T3 cells were cultured for 1 week prior to experiments in supplemented DMEM containing 5% Lipoprotein-Depleted Serum (LDS) (Kalen Biomedical, LLC). To induce ciliation and test for SHH responsiveness, cells were serum starved in 0.5% LDS supplemented DMEM and simultaneously treated with 1 μM U18666A and various concentrations of SHH in the presence or absence of exogenously added cholesterol: methyl-β-cyclodextrin complexes for 24 hours. Due to their inability to survive with prolonged exposure to LDS-containing DMEM, *Lss^-/-^* cells were cultured for 2 days prior to treatment with 1 μM U18666A, SHH, and cholesterol:methyl-β-cyclodextrin complexes for Hedgehog signaling assays. Two or more independent clonal *Dhcr7^-/-^*, *Dhcr24^-/-^* and *Lss^-/-^* cell lines were tested in all assays and gave similar results (**Figs.2B-2C**). Note that the SHH-responsiveness of *Dhcr7^-/-^*, *Dhcr24^-/-^* and *Lss^-/-^* cells was similar to wild-type cells when cultured in media containing lipid-replete serum.

In order to deplete cells of sphingomyelin, myriocin (40 μM, unless otherwise indicated) and fumonisin B1 (40 μM) were added to cells cultured in 10% FBS DMEM for three days prior to HH signaling assays, flow cytometry, or microscopy. Low-dose (50nM) Staurosporine, egg SM: fatty acid free BSA complexes (30 μM), methyl-β-cyclodextrin (300nM), cholesterol: methyl-β-cyclodextrin complexes (300nM), and SHH ligands were all added for 24 hours before analysis unless otherwise indicated. Vehicle controls (such as DMSO for Myriocin and fatty acid free BSA for SM add-back experiments) were used when appropriate. Cholesterol: Methyl-β-cyclodextrin complexes were generated as described previously (Luchetti et al. 2016). Egg sphingomyelin: fatty acid free BSA complexes were made by dissolving in Optimem (Gibco) at a molar ratio of 1000:1 (sphingomyelin:BSA) followed by water-bath sonication.

### Neural Progenitor Differentiation Assays

To assess the effect of myriocin on Hedgehog (HH) signaling in a more physiological, differentiation based assay, we used the HM1 mESC line described previously (Pusapati, Kong, Patel, Gouti, et al. 2018). This cell line harbors the GLI1-Venus and OLIG2-mKate dual reporter system to evaluate the strength of HH signaling output both through *Gli1* target gene induction and the Olig2 differentiation marker for motor neuron progenitors. After growth and maintenance of mESC on feeder cells, the cells were plated on 6-well gelatin-coated CellBIND plates (Corning) at a density of 100,000 cells/well for flow cytometry analysis. Differentiation was carried out in N2B27 media (Dulbecco’s Modified Eagle’s Medium F12 (Gibco) and Neurobasal Medium (Gibco) (1:1 ratio) supplemented with N-2 Supplement (Gibco), B-27 Supplement (Gibco), 1% penicillin/streptomycin (Gemini Bio-Products), 2mM L-glutamine (Gemini Biosciences), 40 mg/mL Bovine Serum Albumin (Sigma), and 55mM 2-mercaptoethanol (Gibco)). Cells were first plated (Day 0) in N2B27 medium supplemented with 10ng/mL bFGF (R&D). One day later (Day 1), either 40 uM myriocin or vehicle control (DMSO) was added to the culture media. On Day 2, media was replaced with bFGF-supplemented media (with or without myriocin) containing 5mM CHIR99021 (Axon). On Day 3, cells were cultured in N2B27 medium containing 100 nM RA (Sigma-Aldrich) (with or without myriocin) and either left untreated (NoSHH) or treated with LoSHH (5 nM) or HiSHH (25 nM). A fresh medium change with the same ingredients was done on Day 4 and Day 5. Finally, on Day 6, cells were washed with PBS and trypsinized for flow cytometry analysis.

### Hedgehog signaling assays

*Gli1* mRNA transcript levels were measured using the Power SYBR Green Cells-to-CT kit (Thermo Fisher Scientific). *Gli1* levels relative to *Gapdh* were calculated using the Delta-Ct method (CT(*Gli1*) - CT(*Gapdh*)). The RT-PCR was carried out using custom primers for *Gli1* (forward primer: 5′-ccaagccaactttatgtcaggg-3′ and reverse primer: 5′-agcccgcttctttgttaatttga-3′), and *Gapdh* (forward primer: 5′-agtggcaaagtggagatt-3′ and reverse primer: 5′-gtggagtcatactggaaca-3′). For analysis of GLI1-GFP in NIH/3T3-CG cells by flow cytometry, cells were harvested by trypsinzation followed by quenching in low (0.5%) serum media at 4°C. Cells were either analyzed by flow cytometry immediately or spun and processed for lipid probe staining. All lipid probes were labeled with red or far-red fluorescent dyes, so GLI-GFP reporter expression and probe staining could be measured simultaneously by two-channel flow cytometry.

### Purification and labeling of lipid probes

Mutant Perfringolysin O (PFO*) was purified as previously described (Akash Das et al. 2014) and covalently labeled with Alexa Fluor 647 following the manufacturer’s instructions. Expression, purification and labeling of OlyA and OlyA_E69A variants containing a single cysteine has been described (Endapally et al. 2019). Both proteins were expressed in Escherichia coli Rosetta(DE3)pLysS cells, purified by metal-affinity and gel-filtration chromatography and finally labelled with Atto-647 maleimide dye following the manufacturer’s instructions (Sigma Aldrich, product #05316).

### Measurement of whole-cell lipid probe staining by flow cytometry

To stain cells with these labeled probes, they were harvested for flow cytometry by trypsinization followed by quenching in ice-cold low (0.5% FBS) serum supplemented DMEM. All subsequent steps were completed on ice. Cells were spun at 1000g and then resuspended in Probe Blocking Buffer (PBB, 1x PBS with 10mg/mL BSA). After 10 minutes in PBB, cells were spun and then resuspended in PBB containing desired probes at the following final concentrations: 5 µg/mL PFO*, 2 µM OlyA and 2 µM OlyA_E69A. Cells were stained for 1 hour and then washed 3 times in PBB before flow cytometry.

### Measurement of ciliary probe staining by microscopy

Probe staining at cilia was carried out using NIH/3T3 or *Ptch^-/-^* cells stably expressing ARL13B-GFP as a cilia marker to avoid the use of detergents for permeabilization. For PFO* probe staining, the coverslips were transferred to an ice-cold metal rack and intact cells were stained in PFO* (at a final concentration of 5 µg/mL) diluted into ice cold low (0.5% FBS) serum supplemented DMEM for 30 minutes. For OlyA and OlyA_E69A probe staining, live cells were stained at room temperature in probe (at a final concentration of 2 µM for both OlyA and OlyA_E69A) diluted in room temperature low (0.5% FBS) serum supplemented DMEM for 10 minutes. After staining, cells were washed with 1x PBS and then immediately fixed in 4% PFA in 1xPBS for 10 minutes. Coverslips were then washed three times with 1x PBS and mounted on glass slides in antifade mountant media (ProLong Diamond Antifade Mountant, Thermo Fisher Scientific) where they cured overnight at room temperature before imaging.

### Thin layer chromatography

Cells grown in the presence or absence of myriocin (40 μM for 3 days) were harvested by washing with 4°C 1x PBS, scraping and spinning at 1000g. Lipids were extracted using chloroform/methanol/water (2:2:1). Lipid extracts were loaded onto a TLC plate (Millipore) along with egg and milk sphingomyelin as lipid standards. The plate was run in a chloroform/acetone/methanol/acetic acid/water (6:8:2:2:1) solvent system, visualized with a 0.03% Coomassie blue G, 100 mM NaCl, 30% methanol solution, and destained with a 100 mM NaCl and 30% methanol solution.

### Analysis of Lipid Probe Staining at Cilia by Immunofluorescence

Images were obtained using a Leica TCS SP5 confocal imaging system containing a 63x oil immersion objective. Quantification of probe staining at cilia was performed using code written in MATLAB R2014b using the following steps. Leica Image Files (LIF) were converted into matrices using the bfmatlab toolbox. Following a max-z-projection, cilia were identified by applying a two-dimensional median filter followed by a high-pass user-defined threshold for signal versus noise to generate a cilia mask in the cilia channel. Any group of contiguous pixels that had signal was labeled as a “potential cilium.” Each potential cilium was then subjected to a series of tests (measuring area, eccentricity, solidity, intensity, and length). If each test was passed, the “true cilium” pixels were then mapped to a matrix containing the pixels in the lipid probe channel. In the lipid probe channel, each focal plane was measured independently to avoid noise caused by probe staining outside of the focal plane of a given cilium. The average intensity of pixels falling within a given cilia mask were measured for each focal plane and these values were recorded in a matrix. In order to measure the local plasma membrane fluorescence around each cilium, the cilia mask was dilated to a user-defined size, and the initial cilia mask was subtracted from the dilated cilia mask in order to create the “plasma membrane mask.” Plasma membrane probe staining was then measured by averaging the pixel intensities within the plasma membrane mask in each focal plane. To generate the final intensity value for each cilium, the focal plane containing the highest average ciliary probe staining was normalized (either by subtraction or division when specified) to the focal plane containing the highest average plasma membrane probe staining. This MATLAB code is available on GitHub (https://github.com/mkinnebr/lipids-HH).

### Statistics

The statistical significance of differences between two groups was determined by an unpaired, nonparametric t-test (Mann-Whitney test). When three or more groups were compared a nonparametric ANOVA (Kruskal-Wallis test) was used. All experiments were repeated at least 3 times.

## Acknowledgements

We thank Kyle Travaglini and Onn Brandman for help with the MATLAB code for automated quantitation of probe fluorescence at primary cilia, Xiaohui Zha and Kevin Courtney for helpful discussions and protocols for sphingomyelin assays, and Suzanne Pfeffer for comments on the manuscript. CS was supported by grants from Cancer Research UK (C20724/A14414 and C20724/A26752) and a European Research Council grant (647278), RR by grants from the National Institutes of Health (GM118082 and GM106078), AR and KAJ by grants from the NIH (HL20948) and Welch Foundation (I-1793), MK and EJI by pre-doctoral fellowships from the National Science Foundation, GL by a pre-doctoral fellowship from the Ford Foundation, GP by a post-doctoral fellowship from the American Heart Association (14POST20370057), and JK by a post-doctoral fellowship from the American Heart Association (19POST34380734) and a K99/R00 award from the NIH (GM13251801).

## Author contributions

RR and AR designed the project. BP analyzed the screen data, with the exception of the KEGG analysis which was performed by MK. BP and GP designed and cloned the CRISPR library focused on lipid-related genes. BP, GP and JK generated the mutant cell library and performed the HiSHH-Bottom10% screen. MK performed the LoSHH-Top5% screen. MK performed the experiments related to cholesterol genes and MK and EJI performed experiments related to sphingomyelin genes. MK, GL and KAJ performed the lipid probe staining experiments. MK and BP designed the computational pipeline for probe quantitation at primary cilia. CS and DFC contributed key conceptual and structural insights and experimental suggestions. RR and MK wrote the paper, with input from all authors.

---

**Supplementary Figure 3 (Related to main Figure 3).**
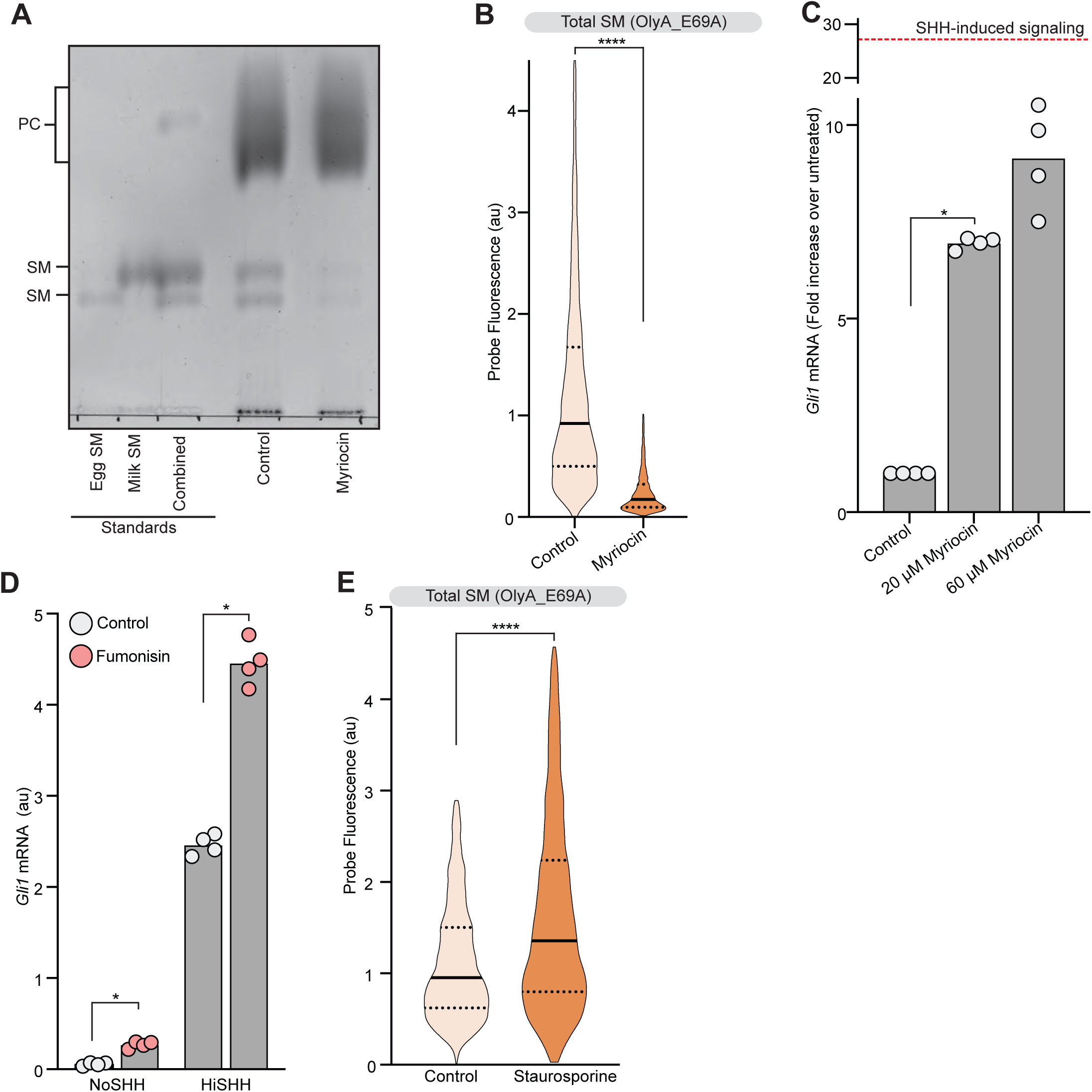
Analysis of sphingomyelin levels and their effect on Hedgehog signaling. (A) Thin Layer Chromatography was used to measure sphingomyelin and phosphatidylcholine (PC) levels in cells treated with myriocin. Purified egg and milk sphingomyelin were used as standards. (B) Flow cytometry was used to measure plasma membrane OlyA_E69A staining in intact cells after myriocin treatment (n> 4000 cells per treatment). (C) HH signaling triggered by myriocin alone in *Smo*^-/-^ MEFs stably expressing wild type SMO. Red bar indicates level of HH signaling induced by HiSHH (25 nM) treatment. (D) HH signaling triggered by Fumonisin B1 in the presence or absence of HiSHH (25 nM). (E) Flow cytometry used to measure plasma membrane OlyA_E69A staining in intact cells after staurosporine treatment (n> 4000 cells per treatment). Statistical significance was determined by the Mann-Whitney test (B); \**p*-value≤ 0.05, \*\*\*\**p*-value≤0.0001.

**Supplementary Figure 4 (Related to Main Figure 4).**
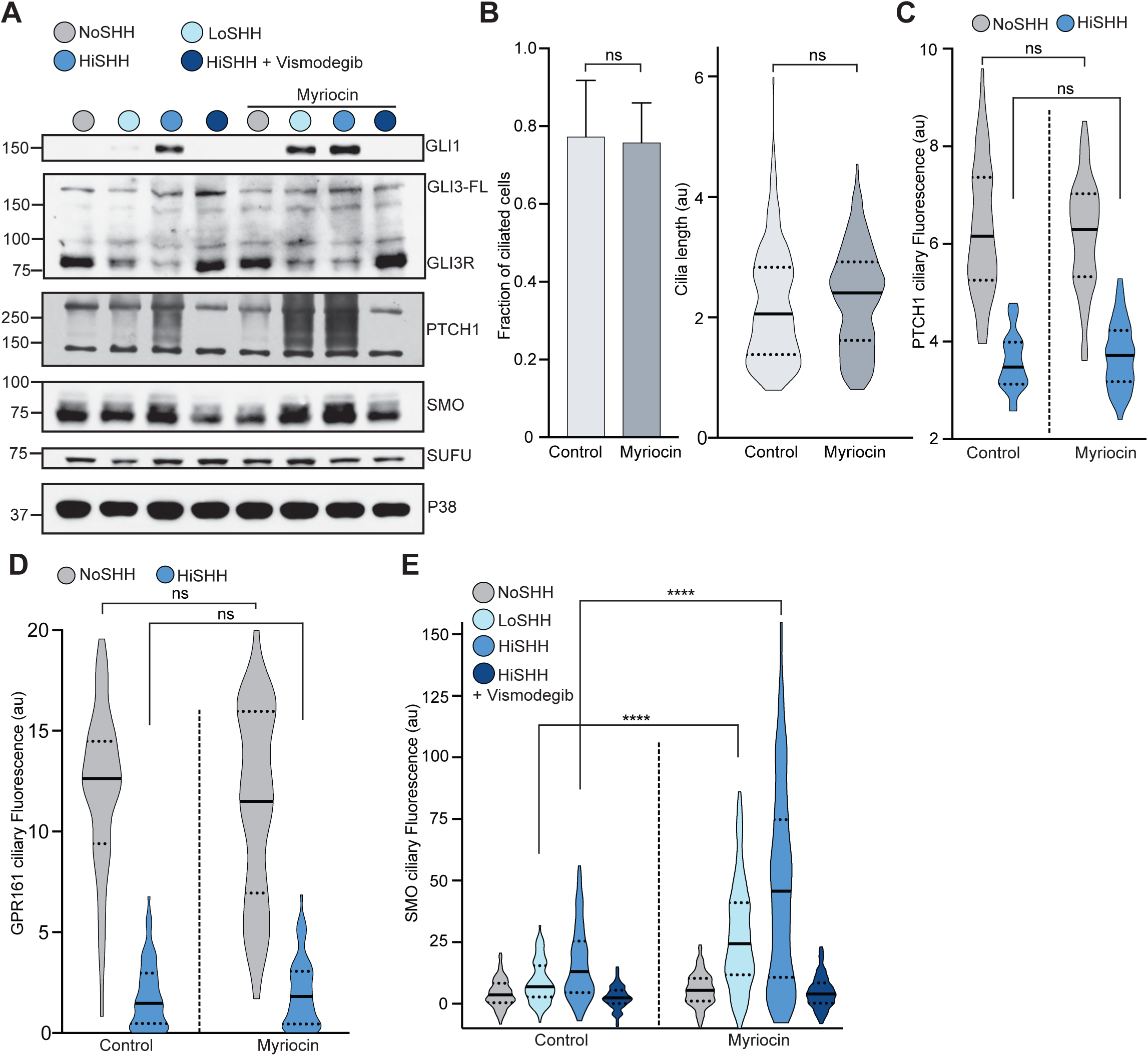
The effect of myriocin treatment on the level of Hedgehog pathway components and ciliary protein trafficking. (A) Western Blot showing levels of HH signaling proteins (SMO, SUFU and GLI3-FL/GLI3R) and HH target genes (GLI1 and PTCH1) after treatment with myriocin and either LoSHH (5 nM), HiSHH (25 nM) or HiSHH with Vismodegib (2.5 uM). (B) Microscopy based analysis showing the fraction of cells containing cilia (calculated as number cilia over number of dapi-stained nuclei, n >100 cilia) and the length of cilia (n = 100) after myriocin treatment. (C and D) Violin plots showing the HiSHH-induced (25 nM) ciliary clearance of endogenous PTCH1 (C, n > 50 cilia per condition) and stably expressed GPR161-YFP (D, n > 35 cilia per condition) in the presence and absence of myriocin. (E) Violin plots showing the SHH-induced ciliary accumulation of endogenous SMO in the presence and absence of myriocin (n > 100 cilia per condition, same SHH concentrations as in (A)). Statistical significance was determined by the Mann-Whitney test (B); \*\*\*\**p*-value≤0.0001, non-significant (ns) *p*-value>0.05.

**Supplementary Figure 6 (Related to Main Figure 6).**
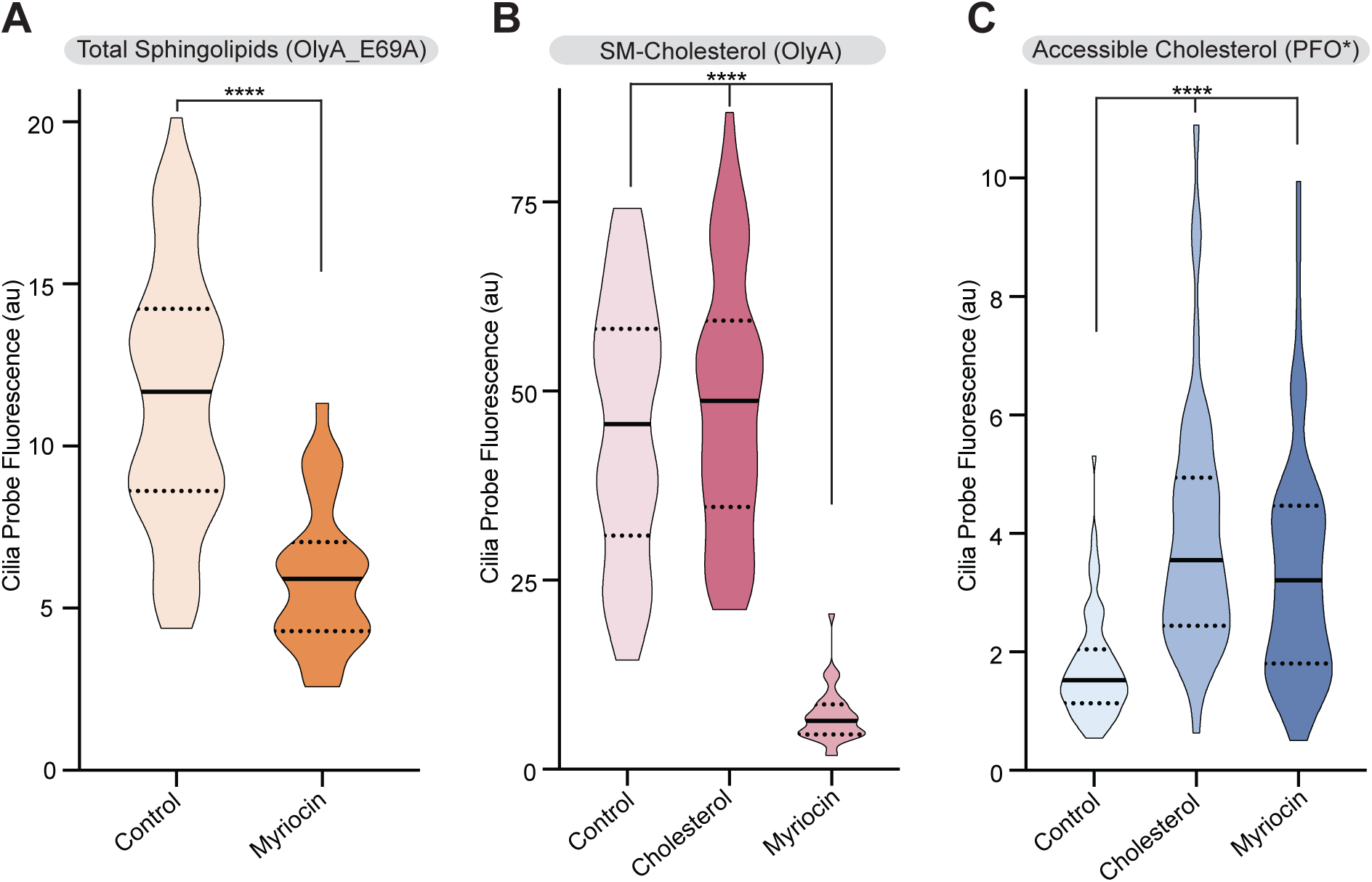
Sphingomyelin depletion increases accessible cholesterol at cilia. (A-C) Ciliary staining intensity for OlyA_E69A, OlyA or PFO* in NIH/3T3 cells left untreated (A) or treated with either myriocin or cholesterol: MβCD complexes (B and C) (n>28 cilia). Statistical significance was determined by the Mann-Whitney test (B); \*\*\*\**p*-value≤0.0001.

**Supplementary Figure 7 (Related to Main Figure 7).**
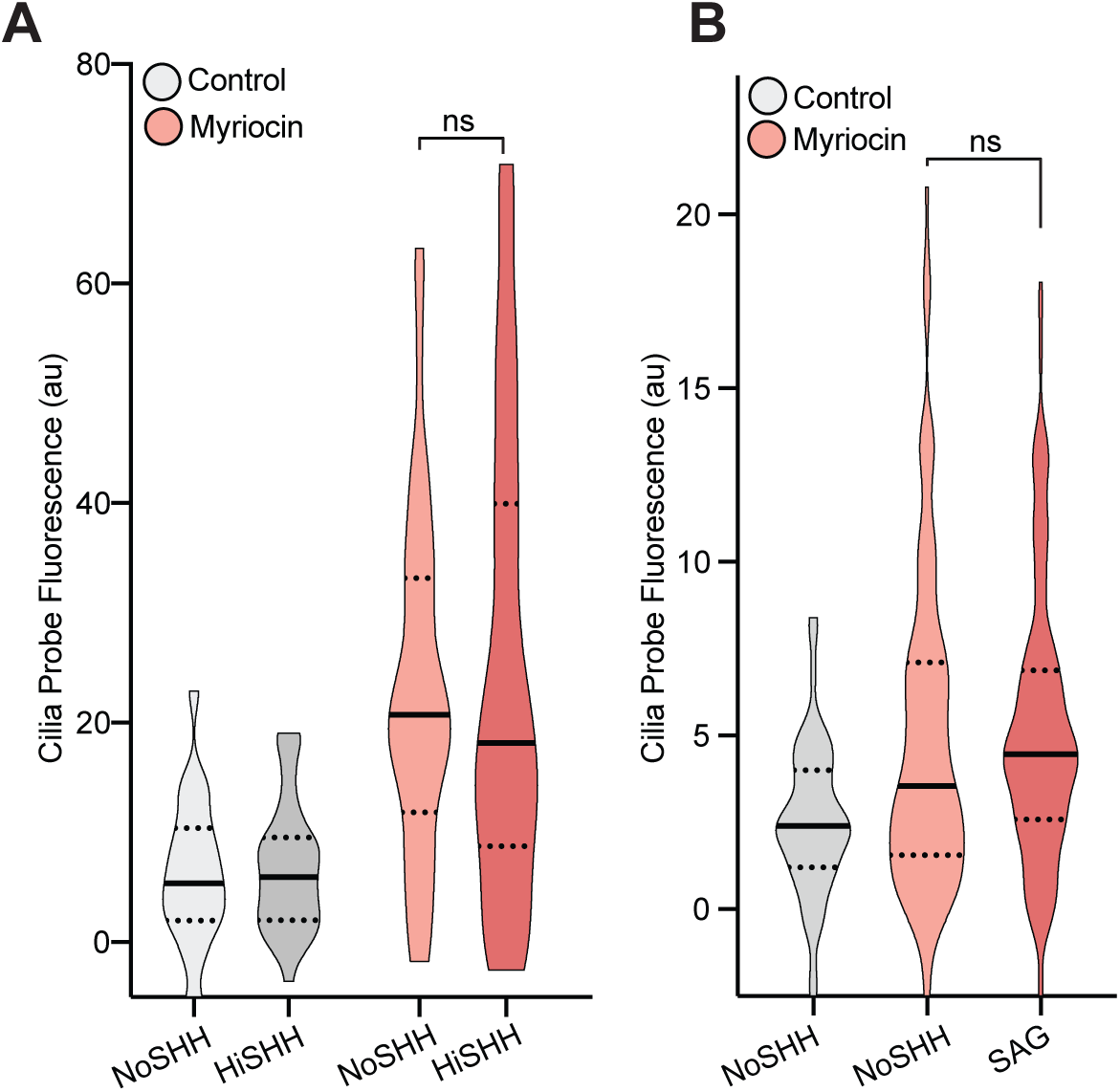
SHH-induced changes in accessible cholesterol levels at cilia depends on PTCH1. (A) PFO* staining at primary cilia in *Ptch1*^-/-^ MEFs in the presence of HiSHH (25 nM, 15 minute treatment) in cells treated with myriocin or left untreated (n>38 cilia). (B) PFO* staining at primary cilia in the presence of SAG (100 nM, 15 minute treatment) in cells treated with myriocin or left untreated (n>50 cilia). Statistical significance was determined by the Mann-Whitney test (B); non-significant (ns) *p*-value>0.05.

## References

Abdel-Khalik, Jonas, Peter J. Crick, Eylan Yutuc, Andrea E. DeBarber, P. Barton Duell, Robert D. Steiner, Ioanna Laina, Yuqin Wang, and William J. Griffiths. 2018. “Identification of 7α, 24-Dihydroxy-3-Oxocholest-4-En-26-Oic and 7α, 25-Dihydroxy-3-Oxocholest-4-En-26-Oic Acids in Human Cerebrospinal Fluid and Plasma.” Biochimie 153: 86–98.

Arensdorf, Angela M., Miriam E. Dillard, Jacob M. Menke, Matthew W. Frank, Charles O. Rock, and Stacey K. Ogden. 2017. “Sonic Hedgehog Activates Phospholipase A2 to Enhance Smoothened Ciliary Translocation.” Cell Reports 19 (10): 2074–87.

Bidet, M., O. Joubert, B. Lacombe, M. Ciantar, R. Nehme, P. Mollat, L. Bretillon, et al. 2011. “The Hedgehog Receptor Patched Is Involved in Cholesterol Transport.” PloS One 6 (9): e23834.

Bielska, Agata A., Paul Schlesinger, Douglas F. Covey, and Daniel S. Ory. 2012. “Oxysterols as Non-Genomic Regulators of Cholesterol Homeostasis.” Trends in Endocrinology and Metabolism: TEM 23 (3): 99–106.

Bijlsma, M. F., C. A. Spek, D. Zivkovic, S. van de Water, F. Rezaee, and M. P. Peppelenbosch. 2006. “Repression of Smoothened by Patched-Dependent (Pro-)Vitamin D3 Secretion.” PLoS Biology 4 (8). http://www.ncbi.nlm.nih.gov/entrez/query.fcgi?cmd=Retrieve&db=PubMed&dopt=Citation&list_uids=16895 439.

Blassberg, R., J. I. Macrae, J. Briscoe, and J. Jacob. 2016. “Reduced Cholesterol Levels Impair Smoothened Activation in Smith-Lemli-Opitz Syndrome.” Human Molecular Genetics 25 (4): 693–705.

Brown, Michael S., Arun Radhakrishnan, and Joseph L. Goldstein. 2018. “Retrospective on Cholesterol Homeostasis: The Central Role of Scap.” Annual Review of Biochemistry 87 (June): 783–807.

Brown, M. S., and J. L. Goldstein. 2009. “Cholesterol Feedback: From Schoenheimer’s Bottle to Scap’s MELADL.” Journal of Lipid Research 50 Suppl (April): S15–27.

Bujold, Mattéa, Akila Gopalakrishnan, Emma Nally, and Kirst King-Jones. 2010. “Nuclear Receptor DHR96 Acts as a Sentinel for Low Cholesterol Concentrations in Drosophila Melanogaster.” Molecular and Cellular Biology 30 (3): 793–805.

Byrne, Eamon Fx, Giovanni Luchetti, Rajat Rohatgi, and Christian Siebold. 2018. “Multiple Ligand Binding Sites Regulate the Hedgehog Signal Transducer Smoothened in Vertebrates.” Current Opinion in Cell Biology 51 (April): 81–88.

Byrne, E. F., R. Sircar, P. S. Miller, G. Hedger, G. Luchetti, S. Nachtergaele, M. D. Tully, et al. 2016. “Structural Basis of Smoothened Regulation by Its Extracellular Domains.” Nature 535 (7613): 517–22.

Carvalho, Maria, Dominik Schwudke, Julio L. Sampaio, Wilhelm Palm, Isabelle Riezman, Gautam Dey, Gagan D. Gupta, et al. 2010. “Survival Strategies of a Sterol Auxotroph.” Development 137 (21): 3675–85.

Chávez, Marcelo, Sabrina Ena, Jacqueline Van Sande, Alban de Kerchove d’Exaerde, Stéphane Schurmans, and Serge N. Schiffmann. 2015. “Modulation of Ciliary Phosphoinositide Content Regulates Trafficking and Sonic Hedgehog Signaling Output.” Developmental Cell 34 (3): 338–50.

Colbeau, A., J. Nachbaur, and P. M. Vignais. 1971. “Enzymac Characterization and Lipid Composition of Rat Liver Subcellular Membranes.” Biochimica et Biophysica Acta (BBA) - Biomembranes. https://doi.org/10.1016/0005-2736(71)90123-4.

Cooper, M. K., C. A. Wassif, P. A. Krakowiak, J. Taipale, R. Gong, R. I. Kelley, F. D. Porter, and P. A. Beachy. 2003. “A Defective Response to Hedgehog Signaling in Disorders of Cholesterol Biosynthesis.” Nature Genetics 33 (4): 508–13.

Corbit, K. C., P. Aanstad, V. Singla, A. R. Norman, D. Y. Stainier, and J. F. Reiter. 2005. “Vertebrate Smoothened Functions at the Primary Cilium.” Nature 437 (7061): 1018–21.

Corcoran, R. B., and M. P. Scott. 2006. “Oxysterols Stimulate Sonic Hedgehog Signal Transduction and Proliferation of Medulloblastoma Cells.” Proceedings of the National Academy of Sciences of the United States of America 103 (22): 8408–13.

Courtney, K. C., W. Pezeshkian, R. Raghupathy, C. Zhang, A. Darbyson, J. H. Ipsen, D. A. Ford, H. Khandelia, J. F. Presley, and X. Zha. 2018. “C24 Sphingolipids Govern the Transbilayer Asymmetry of Cholesterol and Lateral Organization of Model and Live-Cell Plasma Membranes.” Cell Reports 24 (4): 1037–49.

Das, A., J. L. Goldstein, D. D. Anderson, M. S. Brown, and A. Radhakrishnan. 2013. “Use of Mutant 125I-Perfringolysin O to Probe Transport and Organization of Cholesterol in Membranes of Animal Cells.” Proceedings of the National Academy of Sciences of the United States of America 110 (26): 10580–85.

Das, Akash, Michael S. Brown, Donald D. Anderson, Joseph L. Goldstein, and Arun Radhakrishnan. 2014. “Three Pools of Plasma Membrane Cholesterol and Their Relation to Cholesterol Homeostasis.” eLife 3 (June). https://doi.org/10.7554/eLife.02882.

Deshpande, Ishan, Jiahao Liang, Danielle Hedeen, Kelsey J. Roberts, Yunxiao Zhang, Betty Ha, Naomi R. Latorraca, et al. 2019. “Smoothened Stimulation by Membrane Sterols Drives Hedgehog Pathway Activity.” Nature. https://doi.org/10.1038/s41586-019-1355-4.

Dobrosotskaya, I. Y., A. C. Seegmiller, M. S. Brown, J. L. Goldstein, and R. B. Rawson. 2002. “Regulation of SREBP Processing and Membrane Lipid Production by Phospholipids in Drosophila.” Science 296 (5569): 879–83.

Doench, J. G., N. Fusi, M. Sullender, M. Hegde, E. W. Vaimberg, K. F. Donovan, I. Smith, et al. 2016. “Optimized sgRNA Design to Maximize Activity and Minimize off-Target Effects of CRISPR-Cas9.” Nature Biotechnology 34 (2): 184–91.

Dwyer, J. R., N. Sever, M. Carlson, S. F. Nelson, P. A. Beachy, and F. Parhami. 2007. “Oxysterols Are Novel Activators of the Hedgehog Signaling Pathway in Pluripotent Mesenchymal Cells.” The Journal of Biological Chemistry 282 (12): 8959–68.

Endapally, Shreya, Donna Frias, Magdalena Grzemska, Austin Gay, Diana R. Tomchick, and Arun Radhakrishnan. 2019. “Molecular Discrimination between Two Conformations of Sphingomyelin in Plasma Membranes.” Cell 176 (5): 1040–53.e17.

Flanagan, John J., Rodney K. Tweten, Arthur E. Johnson, and Alejandro P. Heuck. 2009. “Cholesterol Exposure at the Membrane Surface Is Necessary and Sufficient to Trigger Perfringolysin O Binding.” Biochemistry 48 (18): 3977–87.

Gale, S. E., E. J. Westover, N. Dudley, K. Krishnan, S. Merlin, D. E. Scherrer, X. Han, et al. 2009. “Side Chain Oxygenated Cholesterol Regulates Cellular Cholesterol Homeostasis through Direct Sterol-Membrane Interactions.” The Journal of Biological Chemistry 284 (3): 1755–64.

Garcia-Gonzalo, Francesc R., Siew C. Phua, Elle C. Roberson, Galo Garcia 3rd, Monika Abedin, Stéphane Schurmans, Takanari Inoue, and Jeremy F. Reiter. 2015. “Phosphoinositides Regulate Ciliary Protein Trafficking to Modulate Hedgehog Signaling.” Developmental Cell 34 (4): 400–409.

Geneva, Ivayla I., Han Yen Tan, and Peter D. Calvert. 2017. “Untangling Ciliary Access and Enrichment of Two Rhodopsin-like Receptors Using Quantitative Fluorescence Microscopy Reveals Cell-Specific Sorting Pathways.” Molecular Biology of the Cell 28 (4): 554–66.

Gijs, Hannah Laura, Nicolas Willemarck, Frank Vanderhoydonc, Niamat Ali Khan, Jonas Dehairs, Rita Derua, Etienne Waelkens, et al. 2015. “Primary Cilium Suppression by SREBP1c Involves Distortion of Vesicular Trafficking by PLA2G3.” Molecular Biology of the Cell 26 (12): 2321–32.

Gong, Xin, Hongwu Qian, Pingping Cao, Xin Zhao, Qiang Zhou, Jianlin Lei, and Nieng Yan. 2018. “Structural Basis for the Recognition of Sonic Hedgehog by Human Patched1.” Science 361 (6402). https://doi.org/10.1126/science.aas8935.

Griffiths, William J., Jonas Abdel-Khalik, Eylan Yutuc, Alwena H. Morgan, Ian Gilmore, Thomas Hearn, and Yuqin Wang. 2017. “Cholesterolomics: An Update.” Analytical Biochemistry 524 (May): 56–67.

Griffiths, William J., Peter J. Crick, Anna Meljon, Spyridon Theofilopoulos, Jonas Abdel-Khalik, Eylan Yutuc, Josie E. Parker, et al. 2019. “Additional Pathways of Sterol Metabolism: Evidence from Analysis of Cyp27a1−/− Mouse Brain and Plasma.” Biochimica et Biophysica Acta (BBA) - Molecular and Cell Biology of Lipids 1864 (2): 191–211.

Griffiths, William J., and Yuqin Wang. 2018. “An Update on Oxysterol Biochemistry: New Discoveries in Lipidomics.” Biochemical and Biophysical Research Communications 504 (3): 617–22.

Griffiths, William J., Eylan Yutuc, Jonas Abdel-Khalik, Peter J. Crick, Thomas Hearn, Alison Dickson, Brian W. Bigger, et al. 2019. “Metabolism of Non-Enzymatically Derived Oxysterols: Clues from Sterol Metabolic Disorders.” Free Radical Biology & Medicine, April. https://doi.org/10.1016/j.freeradbiomed.2019.04.020.

Horvat, Simon, Jim McWhir, and Damjana Rozman. 2011. “Defects in Cholesterol Synthesis Genes in Mouse and in Humans: Lessons for Drug Development and Safer Treatments.” Drug Metabolism Reviews 43 (1): 69–90.

Hotze, Eileen M., Alejandro P. Heuck, Daniel M. Czajkowsky, Zhifeng Shao, Arthur E. Johnson, and Rodney K. Tweten. 2002. “Monomer-Monomer Interactions Drive the Prepore to Pore Conversion of a β-Barrel-Forming Cholesterol-Dependent Cytolysin.” The Journal of Biological Chemistry 277 (13): 11597–605.

Huangfu, D., A. Liu, A. S. Rakeman, N. S. Murcia, L. Niswander, and K. V. Anderson. 2003. “Hedgehog Signalling in the Mouse Requires Intraflagellar Transport Proteins.” Nature 426 (6962): 83–87.

Huang, Pengxiang, Sanduo Zheng, Bradley M. Wierbowski, Youngchang Kim, Daniel Nedelcu, Laura Aravena, Jing Liu, Andrew C. Kruse, and Adrian Salic. 2018. “Structural Basis of Smoothened Activation in Hedgehog Signaling.” Cell 175 (1): 295–97.

Huang, P., D. Nedelcu, M. Watanabe, C. Jao, Y. Kim, J. Liu, and A. Salic. 2016. “Cellular Cholesterol Directly Activates Smoothened in Hedgehog Signaling.” Cell 166 (5): 1176–87 e14.

Infante, Rodney Elwood, and Arun Radhakrishnan. 2017. “Continuous Transport of a Small Fraction of Plasma Membrane Cholesterol to Endoplasmic Reticulum Regulates Total Cellular Cholesterol.” eLife 6 (April). https://doi.org/10.7554/eLife.25466.

Jiang, Kai, Yajuan Liu, Junkai Fan, Jie Zhang, Xiang-An Li, B. Mark Evers, Haining Zhu, and Jianhang Jia. 2016. “PI(4)P Promotes Phosphorylation and Conformational Change of Smoothened through Interaction with Its C-Terminal Tail.” PLoS Biology 14 (2): e1002375.

Joung, Julia, Silvana Konermann, Jonathan S. Gootenberg, Omar O. Abudayyeh, Randall J. Platt, Mark D. Brigham, Neville E. Sanjana, and Feng Zhang. 2017. “Genome-Scale CRISPR-Cas9 Knockout and Transcriptional Activation Screening.” Nature Protocols 12 (4): 828–63.

Khaliullina, H., M. Bilgin, J. L. Sampaio, A. Shevchenko, and S. Eaton. 2015. “Endocannabinoids Are Conserved Inhibitors of the Hedgehog Pathway.” Proceedings of the National Academy of Sciences of the United States of America 112 (11): 3415–20.

Kim, J., J. E. Lee, S. Heynen-Genel, E. Suyama, K. Ono, K. Lee, T. Ideker, P. Aza-Blanc, and J. G. Gleeson. 2010. “Functional Genomic Screen for Modulators of Ciliogenesis and Cilium Length.” Nature 464 (7291): 1048–51.

Kong, Jennifer H., Christian Siebold, and Rajat Rohatgi. 2019. “Biochemical Mechanisms of Vertebrate Hedgehog Signaling.” Development 146 (10). https://doi.org/10.1242/dev.166892.

Lange, Y., H. B. Cutler, and T. L. Steck. 1980. “The Effect of Cholesterol and Other Intercalated Amphipaths on the Contour and Stability of the Isolated Red Cell Membrane.” The Journal of Biological Chemistry 255 (19): 9331–37.

Lange, Y., M. H. Swaisgood, B. V. Ramos, and T. L. Steck. 1989. “Plasma Membranes Contain Half the Phospholipid and 90% of the Cholesterol and Sphingomyelin in Cultured Human Fibroblasts.” The Journal of Biological Chemistry 264 (7): 3786–93.

Lange, Yvonne, Jin Ye, and Theodore L. Steck. 2004. “How Cholesterol Homeostasis Is Regulated by Plasma Membrane Cholesterol in Excess of Phospholipids.” Proceedings of the National Academy of Sciences of the United States of America 101 (32): 11664–67.

Lebensohn, A. M., R. Dubey, L. R. Neitzel, O. Tacchelly-Benites, E. Yang, C. D. Marceau, E. M. Davis, et al. 2016. “Comparative Genetic Screens in Human Cells Reveal New Regulatory Mechanisms in WNT Signaling.” eLife 5. https://doi.org/10.7554/eLife.21459.

“LIPID MAPS Proteome Database.” n.d. https://www.lipidmaps.org/data/proteome/LMPD.php.

Li, W., H. Xu, T. Xiao, L. Cong, M. I. Love, F. Zhang, R. A. Irizarry, J. S. Liu, M. Brown, and X. S. Liu. 2014. “MAGeCK Enables Robust Identification of Essential Genes from Genome-Scale CRISPR/Cas9 Knockout Screens.” Genome Biology 15 (12): 554.

Luchetti, G., R. Sircar, J. H. Kong, S. Nachtergaele, A. Sagner, E. F. Byrne, D. F. Covey, C. Siebold, and R. Rohatgi. 2016. “Cholesterol Activates the G-Protein Coupled Receptor Smoothened to Promote Hedgehog Signaling.” eLife 5. https://doi.org/10.7554/eLife.20304.

Maekawa, Masashi, Minhyoung Lee, Kuiru Wei, Neale D. Ridgway, and Gregory D. Fairn. 2016. “Staurosporines Decrease ORMDL Proteins and Enhance Sphingomyelin Synthesis Resulting in Depletion of Plasmalemmal Phosphatidylserine.” Scientific Reports 6 (November): 35762.

McConnell, Harden M., and Arun Radhakrishnan. 2003. “Condensed Complexes of Cholesterol and Phospholipids.” Biochimica et Biophysica Acta 1610 (2): 159–73.

Mukhopadhyay, S., X. Wen, N. Ratti, A. Loktev, L. Rangell, S. J. Scales, and P. K. Jackson. 2013. “The Ciliary G-Protein-Coupled Receptor Gpr161 Negatively Regulates the Sonic Hedgehog Pathway via cAMP Signaling.” Cell 152 (1-2): 210–23.

Myers, Benjamin R., Lila Neahring, Yunxiao Zhang, Kelsey J. Roberts, and Philip A. Beachy. 2017. “Rapid, Direct Activity Assays for Smoothened Reveal Hedgehog Pathway Regulation by Membrane Cholesterol and Extracellular Sodium.” Proceedings of the National Academy of Sciences of the United States of America 114 (52): E11141–50.

Nachtergaele, S., L. K. Mydock, K. Krishnan, J. Rammohan, P. H. Schlesinger, D. F. Covey, and R. Rohatgi. 2012. “Oxysterols Are Allosteric Activators of the Oncoprotein Smoothened.” Nature Chemical Biology 8 (2): 211–20.

Nachury, Maxence V., and David U. Mick. 2019. “Establishing and Regulating the Composition of Cilia for Signal Transduction.” Nature Reviews. Molecular Cell Biology, April. https://doi.org/10.1038/s41580-019-0116-4.

Ocbina, P. J., and K. V. Anderson. 2008. “Intraflagellar Transport, Cilia, and Mammalian Hedgehog Signaling: Analysis in Mouse Embryonic Fibroblasts.” Developmental Dynamics: An Official Publication of the American Association of Anatomists 237 (8): 2030–38.

Porter, Forbes D., and Gail E. Herman. 2011. “Malformation Syndromes Caused by Disorders of Cholesterol Synthesis.” Journal of Lipid Research 52 (1): 6–34.

Pusapati, Ganesh V., Jennifer H. Kong, Bhaven B. Patel, Mina Gouti, Andreas Sagner, Ria Sircar, Giovanni Luchetti, Philip W. Ingham, James Briscoe, and Rajat Rohatgi. 2018. “G Protein-Coupled Receptors Control the Sensitivity of Cells to the Morphogen Sonic Hedgehog.” Science Signaling 11 (516). https://doi.org/10.1126/scisignal.aao5749.

Pusapati, Ganesh V., Jennifer H. Kong, Bhaven B. Patel, Arunkumar Krishnan, Andreas Sagner, Maia Kinnebrew, James Briscoe, L. Aravind, and Rajat Rohatgi. 2018. “CRISPR Screens Uncover Genes That Regulate Target Cell Sensitivity to the Morphogen Sonic Hedgehog.” Developmental Cell 44 (1): 113–29.e8.

Qian, Hongwu, Pingping Cao, Miaohui Hu, Shuai Gao, Nieng Yan, and Xin Gong. 2019. “Inhibition of Tetrameric Patched1 by Sonic Hedgehog through an Asymmetric Paradigm.” Nature Communications 10 (1): 2320.

Qi, Chao, Giulio Di Minin, Irene Vercellino, Anton Wutz, and Volodymyr M. Korkhov. 2018. “Structural Basis of Sterol Recognition by Human Hedgehog Receptor PTCH1.” bioRxiv. https://doi.org/10.1101/508325.

Qi, Xiaofeng, Philip Schmiege, Elias Coutavas, and Xiaochun Li. 2018. “Two Patched Molecules Engage Distinct Sites on Hedgehog Yielding a Signaling-Competent Complex.” Science 362 (6410). https://doi.org/10.1126/science.aas8843.

Radhakrishnan, A., T. G. Anderson, and H. M. McConnell. 2000. “Condensed Complexes, Rafts, and the Chemical Activity of Cholesterol in Membranes.” Proceedings of the National Academy of Sciences of the United States of America 97 (23): 12422–27.

Radhakrishnan, Arun, Joseph L. Goldstein, Jeffrey G. McDonald, and Michael S. Brown. 2008. “Switch-like Control of SREBP-2 Transport Triggered by Small Changes in ER Cholesterol: A Delicate Balance.” Cell Metabolism 8 (6): 512–21.

Raleigh, David R., Navdar Sever, Pervinder K. Choksi, Monika Abedin Sigg, Kelly M. Hines, Bonne M. Thompson, Daniel Elnatan, et al. 2018. “Cilia-Associated Oxysterols Activate Smoothened.” Molecular Cell 72 (2): 316–27.e5.

Regard, J. B., D. Malhotra, J. Gvozdenovic-Jeremic, M. Josey, M. Chen, L. S. Weinstein, J. Lu, E. M. Shore, F. S. Kaplan, and Y. Yang. 2013. “Activation of Hedgehog Signaling by Loss of GNAS Causes Heterotopic Ossification.” Nature Medicine 19 (11): 1505–12.

Rietveld, Anton, Stephanie Neutz, Kai Simons, and Suzanne Eaton. 1999. “Association of Sterol- and Glycosylphosphatidylinositol-Linked Proteins withDrosophilaRaft Lipid Microdomains.” Journal of Biological Chemistry. https://doi.org/10.1074/jbc.274.17.12049.

Rohatgi, R., L. Milenkovic, R. B. Corcoran, and M. P. Scott. 2009. “Hedgehog Signal Transduction by Smoothened: Pharmacologic Evidence for a 2-Step Activation Process.” Proceedings of the National Academy of Sciences of the United States of America 106 (9): 3196–3201.

Rohatgi, R., L. Milenkovic, and M. P. Scott. 2007. “Patched1 Regulates Hedgehog Signaling at the Primary Cilium.” Science 317 (5836): 372–76.

Sanjana, Neville E., Ophir Shalem, and Feng Zhang. 2014. “Improved Vectors and Genome-Wide Libraries for CRISPR Screening.” Nature Methods 11 (8): 783–84.

Scheek, S., M. S. Brown, and J. L. Goldstein. 1997. “Sphingomyelin Depletion in Cultured Cells Blocks Proteolysis of Sterol Regulatory Element Binding Proteins at Site 1.” Proceedings of the National Academy of Sciences of the United States of America 94 (21): 11179–83.

Sever, Navdar, Randall K. Mann, Libin Xu, William J. Snell, Carmen I. Hernandez-Lara, Ned A. Porter, and Philip A. Beachy. 2016. “Endogenous B-Ring Oxysterols Inhibit the Hedgehog Component Smoothened in a Manner Distinct from Cyclopamine or Side-Chain Oxysterols.” Proceedings of the National Academy of Sciences. https://doi.org/10.1073/pnas.1604984113.

Sharpe, H. J., W. Wang, R. N. Hannoush, and F. J. de Sauvage. 2015. “Regulation of the Oncoprotein Smoothened by Small Molecules.” Nature Chemical Biology 11 (4): 246–55.

Sharpe, Laura J., and Andrew J. Brown. 2013. “Controlling Cholesterol Synthesis beyond 3-Hydroxy-3-Methylglutaryl-CoA Reductase (HMGCR).” The Journal of Biological Chemistry 288 (26): 18707–15.

Simons, K., and E. Ikonen. 1997. “Functional Rafts in Cell Membranes.” Nature 387 (6633): 569–72.

Simons, K., and E. Ikonen. 2000. “How Cells Handle Cholesterol.” Science 290 (5497): 1721–26.

Skočaj, Matej, Nataša Resnik, Maja Grundner, Katja Ota, Nejc Rojko, Vesna Hodnik, Gregor Anderluh, et al. 2014. “Tracking Cholesterol/sphingomyelin-Rich Membrane Domains with the Ostreolysin A-mCherry Protein.” PloS One 9 (3): e92783.

Slotte, J. P., and E. L. Bierman. 1988. “Depletion of Plasma-Membrane Sphingomyelin Rapidly Alters the Distribution of Cholesterol between Plasma Membranes and Intracellular Cholesterol Pools in Cultured Fibroblasts.” Biochemical Journal 250 (3): 653–58.

Sokolov, Anna, and Arun Radhakrishnan. 2010. “Accessibility of Cholesterol in Endoplasmic Reticulum Membranes and Activation of SREBP-2 Switch Abruptly at a Common Cholesterol Threshold.” The Journal of Biological Chemistry 285 (38): 29480–90.

Steck, Theodore L., and Yvonne Lange. 2010. “Cell Cholesterol Homeostasis: Mediation by Active Cholesterol.” Trends in Cell Biology 20 (11): 680–87.

Steck, Theodore L., and Yvonne Lange. 2018. “Transverse Distribution of Plasma Membrane Bilayer Cholesterol: Picking Sides.” Traffic 19 (10): 750–60.

Tafesse, Fikadu G., Ali Rashidfarrokhi, Florian I. Schmidt, Elizaveta Freinkman, Stephanie Dougan, Michael Dougan, Alexandre Esteban, Takeshi Maruyama, Karin Strijbis, and Hidde L. Ploegh. 2015. “Disruption of Sphingolipid Biosynthesis Blocks Phagocytosis of Candida Albicans.” PLoS Pathogens 11 (10): e1005188.

Tafesse, Fikadu G., Sumana Sanyal, Joseph Ashour, Carla P. Guimaraes, Martin Hermansson, Pentti Somerharju, and Hidde L. Ploegh. 2013. “Intact Sphingomyelin Biosynthetic Pathway Is Essential for Intracellular Transport of Influenza Virus Glycoproteins.” Proceedings of the National Academy of Sciences of the United States of America 110 (16): 6406–11.

Touster, O., N. N. Aronson Jr, J. T. Dulaney, and H. Hendrickson. 1970. “Isolation of Rat Liver Plasma Membranes. Use of Nucleotide Pyrophosphatase and Phosphodiesterase I as Marker Enzymes.” The Journal of Cell Biology 47 (3): 604–18.

Vinci, Giovanna, Xuhua Xia, and Reiner A. Veitia. 2008. “Preservation of Genes Involved in Sterol Metabolism in Cholesterol Auxotrophs: Facts and Hypotheses.” PloS One 3 (8): e2883.

Zhang, Yunxiao, David P. Bulkley, Yao Xin, Kelsey J. Roberts, Daniel E. Asarnow, Ashutosh Sharma, Benjamin R. Myers, Wonhwa Cho, Yifan Cheng, and Philip A. Beachy. 2018. “Structural Basis for Cholesterol Transport-like Activity of the Hedgehog Receptor Patched.” Cell, November. https://doi.org/10.1016/j.cell.2018.10.026.

